# Influence of vegetation structure on lemur recolonization of post-fire habitats in northwestern Madagascar

**DOI:** 10.1101/2025.11.26.690642

**Authors:** Naina Ratsimba Rabemananjara, Misa Rasolozaka, Marie Odile Ravolanirina, Rogula Marivola, Seheno Harilala Randriamiarantsoa, Romule Rakotondravony, Hanta Razafindraibe, Dominik Schüßler, Ute Radespiel

## Abstract

Habitat quality is a key determinant of wildlife distribution and persistence. Any disturbance such as fire degrades forest habitats by increasing canopy openness, reducing tree basal area, and eliminating large, cavity-bearing trees providing shelter for many species. In Madagascar, fire has become a prominent disturbance, yet the mechanisms linking fire disturbance to lemur habitat occupation remain poorly understood. Using Generalized Linear Mixed Models, we assessed the effects of forest humidity, temperature, vegetation structural gradients, and tree species richness on lemur occurrence, and evaluated the impact of fire regime parameters on significant abiotic and structural vegetation gradients. Lemur responses varied by body size and ecological specialization. *Eulemur fulvus* most likely occurred in humid habitats with high floristic diversity. The latter also explained best the presence of *Avahi occidentalis* and *Cheirogaleus medius,* while *Lepilemur edwardsi* was rather connected to dense stands of overstory trees. Small-bodied *Microcebus* species displayed a higher structural and floristic tolerance. Tree species richness declined sharply in burnt areas and was highest in unburnt forest. Fires impacted vegetation structural gradients, increasing openness and understory density, while old burnt and unburnt forests maintained complex vertical layering. Tree species richness emerged as the strongest predictor of lemur species richness. These results indicate that fire primarily affects lemurs through floristic and structural pathways rather than microclimatic shifts, emphasizing the need to preserve intact forest refugia with mature canopies and species-rich tree communities to sustain lemur diversity under increasing fire pressure in dry forest landscapes of Madagascar.

**Summary:**

- Forest fires alter lemur habitats by reducing tree species richness and mature tree density.
- *Eulemur fulvus, Avahi occidentalis* and *Cheirogaleus medius* preferentially occupied floristically diverse forests, suggesting the relevance of a diverse spectrum of food resources for species of divergent body size.
- The medium-sized *Lepilemur edwardsi* was sensitive to fire-induced loss of canopy integrity, limiting their persistence in burnt forests, while small-bodied mouse lemurs (*Microcebus ravelobensis and M. murinus*) exhibited high resilience to fire-modulated habitat changes.
- Lemur species richness was strongly linked to tree species diversity, underscoring the role of floristic richness in post-fire lemur and habitat recovery.

## 1 INTRODUCTION

Habitat quality is a primary driver of wildlife distribution, persistence, and ecological interactions across ecosystems (Flesch, 2017). Structural attributes such as canopy architecture, tree density, and understory complexity can shape resource availability, movement pathways, and predator avoidance (Ishii et al., 2004; McLean et al., 2016). Greater structural heterogeneity generally supports higher biodiversity by creating a wider array of ecological niches (LaRue et al., 2019).

For arboreal mammals, canopy height, connectivity, and substrate diversity are especially critical (Cudney-Valenzuela et al., 2021). Dense, connected canopies enhance safe movement, reduce predation, and promote species richness, conditions typical of old-growth forests with high structural complexity (Cudney-Valenzuela et al., 2022; Lindenmayer et al., 2017). Conversely, logging, fire, and fragmentation increase canopy openness, reduce basal area, and eliminate large trees that provide cavities and therefore shelter, leading to declines in arboreal mammal diversity and abundance, particularly among rare and specialist species (Cudney-Valenzuela et al., 2021; Lindenmayer et al., 2017; Rodríguez-Gómez & Fontúrbel, 2020).

Primates are among the most affected groups (Estrada & Garber, 2022). Loss of canopy cover and vertical complexity constrains their movement and foraging, driving consistent declines in abundance and richness as forest cover decreases (Gestich et al., 2022). Even partial structural degradation can diminish habitat quality and reduce the capacity of forests to sustain primate populations (Harcourt & Doherty, 2005). Matrix composition also influences primate persistence: open or fire-degraded landscapes constrain dispersal and increase population isolation (Fahrig, 2017). Habitat dependence often scales with body size, as large-bodied primates rely more heavily on canopy connectivity and complex structure for locomotion and predator avoidance, whereas smaller species can persist in more disturbed or fragmented habitats (Cant, 1992; Johns & Skorupa, 1987).

Fire is one of the most pervasive ecological forces shaping tropical landscapes. While natural fires occur regularly in savannah, grasslands, boreal forest, and in regions with Mediterranean climates (Keeley et al., 2011; Leys et al., 2018), they have been historically rare in humid tropical forests due to high moisture levels (Bond & Keeley, 2005). However, human activities such as shifting cultivation and land clearing have increased fire frequency and intensity across tropical regions (Cochrane, 2003). Fire removes biomass, opens the canopy, and alters microclimates, often favoring pioneer or fire-tolerant vegetation over mature forest species (Fernandes-Carvalho-De-Andrade & Brazil, 2021; Moser et al., 2010). Repeated burning can drive long-term vegetation shifts toward grass-dominated states, challenging forest resilience and regeneration potential (Cochrane & Laurance, 2002).

Beyond immediate mortality, fire influences fauna primarily through long-term changes in habitat structure, resource availability, and local microclimate (Lyon et al., 2012). In Indian dry forests, for example, fire compels animals to forage in open habitats with reduced resources and higher predation risk (Johnsingh, 1986). In this context, the severity and recurrence of fire events are particularly important, as they determine whether ecosystems retain resilience for natural recovery or cross tipping points towards permanently degraded alternative states (González et al., 2022).

Madagascar is one of the world’s most distinctive biodiversity hotspots, characterized by exceptional endemism with more than 90% of its fauna and flora being found nowhere else on Earth (Goodman & Benstead, 2005). Lemurs, the endemic primate clade of Madagascar, comprise more than 100 species which vary remarkably in body size, ecology, and life history strategies (Mittermeier et al., 2023). Lemurs inhabit nearly all forested habitats from humid rainforests to dry deciduous and spiny forests, but this diversity is increasingly threatened by deforestation, fragmentation, and fires (Mittermeier et al., 2023).

Fire is a dominant ecological and anthropogenic force in Madagascar, historically shaping vegetation and now contributing substantially to forest loss (Phelps et al., 2022). Although some grassy biomes have long been maintained by natural fire regimes, most contemporary fires are human induced, associated with shifting cultivation (tavy), pasture creation, and hunting (Kull, 2004). Recent satellite data revealed widespread fire anomalies across the island, challenging global assumptions about degradation dynamics (Joseph et al., 2023). The western dry deciduous forests are especially fire-prone due to seasonal drought and flammable vegetation, making them one of the most threatened ecosystems worldwide (Nolan et al., 2020; Phelps et al., 2022). In general, recurrent fires can reduce canopy height, increase gap frequency, and promote fire-tolerant shrubs and grasses. Over time, this process impedes regeneration and can permanently convert forests into degraded shrublands or grasslands with diminished capacity to support native fauna (Allerton et al., 2025; Harrison et al., 2024; Pereira et al., 2024). Despite these potential threats, the ecological consequences of fires for Madagascar’s fauna remain poorly understood.

Ankarafantsika National Park (ANP) in northwestern Madagascar represents a key remnant of dry deciduous forest and an ideal setting to study fire–fauna interactions (Alonso et al., 2002). The National Park history of recurrent wildfires (Rasolozaka et al., 2025) has produced a mosaic of forest patches at varying successional stages adjacent to intact primary forest (Gautier et al., 2018). This landscape offers a comparative framework to study lemur communities across a gradient of disturbance. ANP supports a community of eight lemur species: three small-sized, nocturnal species (*Microcebus murinus*, *M. ravelobensis*, *Cheirogaleus medius)*, the medium-sized *Lepilemur edwardsi* and *Avahi occidentalis*, two large-bodied cathemeral species (*Eulemur fulvus, E. mongoz)*, and the large diurnal *Propithecus coquereli* (Mittermeier et al., 2023).

A prior study in ANP showed show that lemur species richness is highest in unburnt forests (Rabemananjara et al., 2025). Large-bodied species such as *Eulemur fulvus*, *E. mongoz*, and *P. coquereli* are largely restricted to unburnt or long-unburnt habitats. Medium-sized species like *Avahi occidentalis* and *Lepilemur edwardsi* occur at low densities in recently burned forests but are absent from frequently burned areas. In contrast, smaller species exhibit greater flexibility: *Microcebus ravelobensis* and *M. murinus* persist even in recently burned sites (Rabemananjara et al., 2025). That study suggested that body size, habitual locomotion, the available of shelter, and therefore essentially habitat structure may be a central factor mediating the species-specific sensitivity to fire, with large-bodied canopy specialists being most vulnerable and small-bodied generalists displaying higher resilience (Rabemananjara et al., 2025).

The present study addresses this hypothesis by examining how variations in vegetation structure and abiotic conditions shape the presence of lemur species of different body sizes in post-fire habitats in northwestern Madagascar. Understanding these relationships is crucial for predicting ecosystem recovery trajectories and developing evidence-based conservation strategies for fire-prone landscapes. Specifically, we pursue two objectives: (1) evaluate whether variations in vegetation structure and abiotic conditions affect the presence of large-, medium-, and small-bodied lemur species, and (2) assess whether these structural and abiotic elements differ depending on the underlying fire history. Based on ecological theory, empirical evidence from other tropical regions, and preliminary observations from Madagascar, we propose the following two hypotheses:

**H1:** Lemur presence will be positively associated with greater structural complexity (e.g., canopy cover, tree species diversity, vertical stratification) and undisturbed abiotic conditions, with large-bodied species showing the strongest dependence on intact canopy connectivity and structural heterogeneity, and small-bodied, generalist species being most tolerant of fire-modulated changes.

**H2:** Those vegetation parameters that influence lemur presence, are impacted by the fire status (burnt/unburnt), fire severity, fire frequency, and the recency of the last fire. Thus, fire-related structural and floristic simplification can explain the species-specific vulnerabilities of lemurs to fires at least partially.

## 2 METHODS

### 2.1 Ethics statement

This study followed the American Society of Primatologists’ Principles for the Ethical Treatment of Non-Human Primates. All fieldwork protocols were reviewed and approved by the Institute of Zoology, University of Veterinary Medicine Hannover, Germany. The research complied with the legal regulations of Madagascar and received authorization from the Ministère de l’Environnement et du Développement Durable (MEDD), the Direction Régionale de l’Environnement (DREDD) Boeny Betsiboka, and Madagascar National Parks (permit numbers: № 275/22/MEDD/SG/DGGE/DAPRNE/SCBE.Re, № 070/23/MEDD/SG/DGGE/DAPRNE/SCBE.Re, and № 263/23/MEDD/SG/DGGE/DAPRNE/SCB E.Re).

### 2.2 Study sites and ecological context

Our study was conducted in Ankarafantsika National Park (ANP), the largest remaining protected area of dry deciduous forest in northwestern Madagascar (−16°08’60”S, 46°57’0”E), covering 1,350 km² (Figure S1). The park’s topography includes river valleys at ∼50 m above sea level (a.s.l.), slopes, and a calcitic plateau that rises southward to ∼350 m a.s.l., forming cliffs in eastern and southern areas (Alonso et al., 2002). Vegetation consists of seasonally dry forests with a mosaic of dry deciduous forests, dry thickets at higher elevations, moist riverine forests around lakes and upstream valleys, and *Raphia* swamp forests in downstream valleys (Goodman et al., 2021). Forest cover occurs across plateaus, slopes, and valleys. Rainfall averages 1,000–1,500 mm annually, concentrated between November and April, while dry season from May to October is almost rainless (Alonso et al., 2002). ANP contains a rich and diverse flora with over 287 species of woody plants of which 92% are endemic to Madagascar (Alonso et al. 2002). The park is also a key refuge of eight lemur species, two of which being Critically Endangered, the Coquerel’s sifaka (*Propithecus coquereli)* and Mongoose lemurs *(Eulemur mongoz)* (Schwitzer et al., 2013).

This study was conducted in the same 18 study sites that were already used for our previous study, representing different fire histories within ANP (Rasolozaka et al., 2025). Briefly, Landsat imagery (30 × 30 m resolution) from 1988–2023 was used to classify pixels as burnt or unburnt for each year based on the Normalized Burn Ratio (NBR) by subtracting pre- and post-fire NBR values (dNBR, (Key & Benson, 2006), and by ground-truthing the inferred fire history at each site. Fire severity was quantified with the same resolution for all sites and years using the dNBR (Keeley, 2009). Sites were chosen to include contrasts between long-unburnt areas and adjacent burnt areas with varying fire frequencies and times since last fire, ensuring a mixed design while controlling for ecological context (Rasolozaka et al., 2025). Consequently, transects were subdivided into partitions based on distinct fire histories. Fieldwork was conducted between September and December 2022 (late dry to early rainy season) and between May and November 2023 (dry season). Transect partitions differed in time since the last fire (1 to >35 years), fire frequency (1–7 events over 35 years), minimum fire intervals (1–31 years), and maximum burn severity.

### 2.3 Collection of vegetation and abiotic data

At each site, we established a 1.2 km transect flagged and GPS-mapped at 10 m intervals (Garmin GPSMAP 64). This transect consisted of two 500 m sections extending perpendicularly from the fire edge into burnt and unburnt forest, with the unburnt line offset by an additional 200 m section placed parallel to the fire edge within 50 m inside the burnt zone, capturing the transition between forest types (Figure 1). Data collection was conducted across all three transect sections.

**FIGURE 1.**
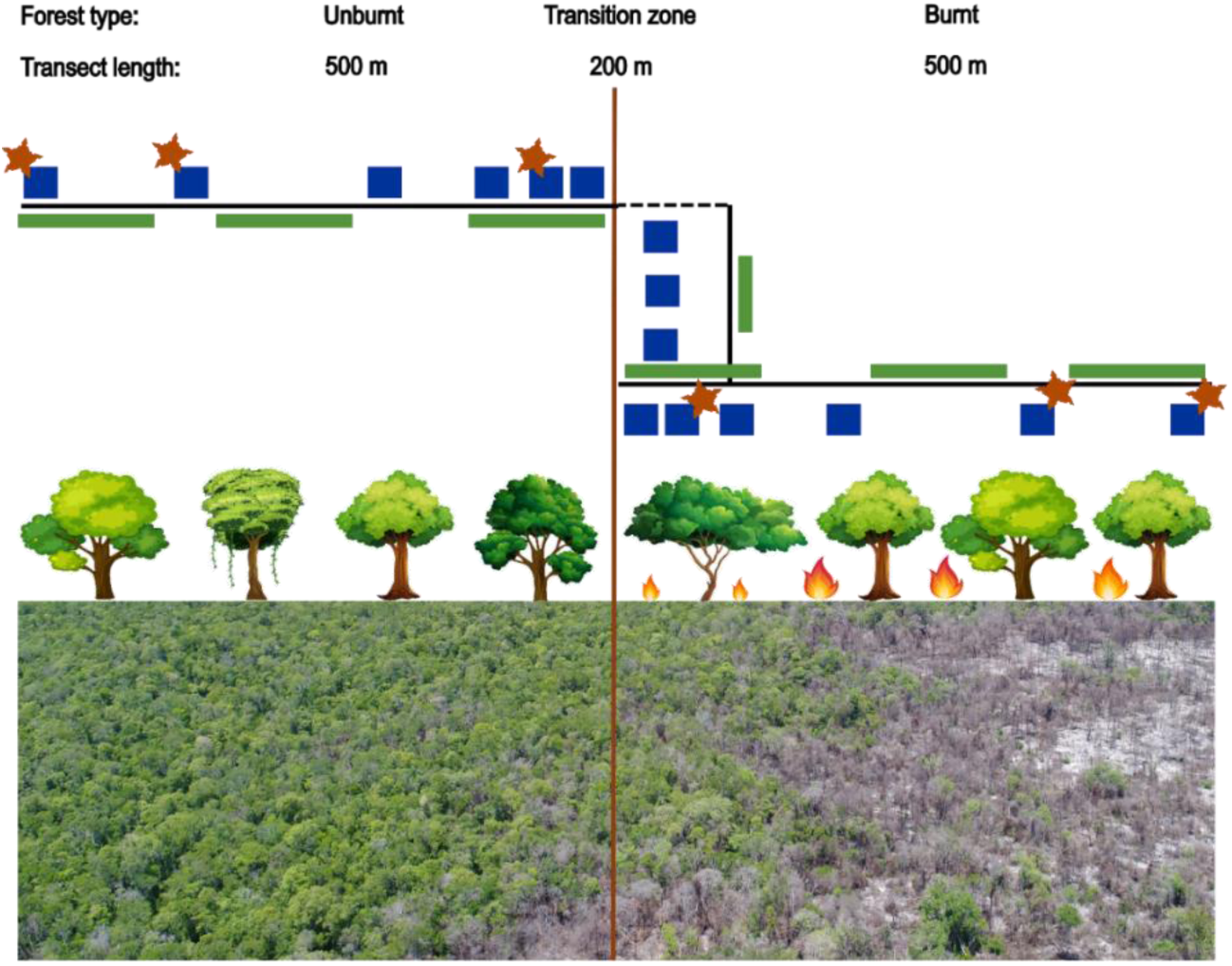
Standardized transect design for the field work. One burnt transect (perpendicular to the fire edge) is paired with an unburnt transect (perpendicular to the fire edge), both being connected by a 200 m long intermediate transition zone transect (50 m inside the burnt forest). Blue squares indicate structural vegetation plots, green lines represent floristic plots, and red stars show locations of abiotic data loggers. The solid black line marks transect lines, and the dashed line indicates a connecting path. Below: Drone image taken a few months after the 2021 fire near the research station of Ampijoroa in ANP illustrating the spatial setting of the transects regarding the fire border. Photograph taken by Hiroki Sato.

Seven rectangular plots (3 m × 50 m) were placed along the 1.2 km transect at each site to capture the floristic variation across habitats with different fire exposure, with three plots located in unburnt forest and four in burnt forest, including one within the transition zone at the fire edge where burnt and unburnt vegetation meet (Figure 1). Within each plot, we recorded all trees with a diameter at breast height (DBH) > 7 cm, identified them to species level whenever possible, and recorded and identified all shrubs taller than 1 m. For each plot, we quantified tree density as the number of individuals and tree species richness as the total number of distinct species.

In addition, 15 structural vegetation plots (5m x 5m) were established along each transect. While six plots were located in the unburnt, nine were placed in the burnt forest, including three positioned in the transition zone (Figure 1). We counted all trees within each plot and categorized them into DBH classes (small: 0.25-5.0 cm; medium: 5.1-10.0 cm; large: >10.1 cm) and height classes (small: 2.5-5.0 m; medium: 5.1-7.50 m; tall: 7.51-10.0 m; very tall: >10.1 m). Finally, forest stratum cover was estimated in these plots for the lower (2.5-5.0 m), medium (5.0–10.0 m), and upper stratum (>10.0 m) by using a modified Londo scale approach with 5% intervals (Londo, 1976).

Abiotic conditions (average relative humidity at 7 a.m. and maximum daily temperature) were measured using automated data loggers (GSP-6, Elitech) placed along transects, with three located in the unburnt forest, three in the burnt forest, and one in the transition zone (Figure 1). We extracted the mean value recorded from all data loggers situated in each transect partition per site during the respective study period.

### 2.4 Determination of lemur species presence

We conducted three diurnal and three nocturnal systematic surveys along each 1.2 km transect to assess the presence of lemur species along its different partitions (Peres, 1999). Four observers walked each transect quietly at ∼0.5 km/h (Rakotondravony & Radespiel, 2007), scanning continuously to detect animals on both sides. Surveys were carried out in the morning (06h30–08h30) and in the evening (18h15–20h30) on three consecutive days. Morning surveys targeted diurnal species, while evening surveys targeted nocturnal species, with headlamps being used to detect eye-shine. Cathemeral species, being active both day and night, were noted whenever encountered. For each detection, we recorded species identity, presence, and the position along the transect to the nearest 10 m.

## 3 STATISTICAL ANALYSIS

Because of the small number of observations, statistical models could not be fitted for *Propithecus coquereli* and *Eulemur mongoz*. *Cheirogaleus medius* undergoes hibernation during much of the dry season (May–September) (Müller & Thalmann, 2002) and was consequently included in the analysis only for the 11 sites surveyed from 5 September onwards in both years. All species-specific models and the those for lemur species richness were fitted in R (R Core Team, 2025) via the RStudio interface (Posit team, 2025).

### 3.1 Modelling the effects of vegetation structure and abiotic factors on lemur presence and lemur species richness

Before fitting models, we tested for correlation among the structural variables using Pearson’s correlation (“stats” package). Since height class counts and the DBH class counts were strongly correlated (r ≥ 0.7, Figure S2), we chose the height class counts to represent them both in the principal component analysis (PCA) analysis. We used a PCA to reduce multicollinearity and to condense the eight parameters included in the vegetation data set (four height classes, tree density, three forest stratum cover estimates). The PCA involved standardizing data, calculating the covariance matrix, and extracting Eigenvalues > 1 to identify principal components (PCs) used for downstream analyses (Table S1). Vegetation variables with loadings > 0.3 were considered important (Table S1) (Hair et al. 2019).

We fitted Generalized Linear Mixed Models (GLMMs) with the “glmmTMB” package (Brooks et al., 2017) to test how vegetation structure (PCs), abiotic factors (humidity, maximum temperature), and tree species richness (species per plot) impacted the presence/absence of each lemur species and lemur species richness (response variables), respectively. Based on data structure and distribution of the response variable, we applied binomial model families. Model fit was evaluated with the “DHARMa” package (Hartig & Hartig, 2022) which included testing for residual distribution, over-/underdispersion, outliers, and zero inflation. If required, we included ziformula() and dispformula() to address zero inflation and dispersion issues, respectively (Brooks et al., 2017). The inclusion of both month and site as a random effect caused convergence problems; therefore, site was excluded from the final model.

We used an information-theoretic approach for model selection with the dredge() function in the “MuMIn” package (Bartoń, 2022). This generates all possible subsets of the global model and ranks them by AICc. Models within ΔAICc ≤ 2 were considered well supported (Burnham & Anderson, 2002) and retained for further analysis (Table S2). To account for model uncertainty, we applied conditional model averaging with the model.avg() function, restricted to the well supported set of models (Burnham & Anderson, 2003; Dormann et al., 2018; Le & Clarke, 2022). This method averages estimates only across models where a variable appears, reducing dilution from absent variables (Symonds & Moussalli, 2011), and is preferred over selecting a single best model because it incorporates uncertainty from multiple plausible models. Model-averaged coefficients, standard errors, and 95% confidence intervals were calculated, and each variables’ importance (RI) was assessed from the summed Akaike weights across the top model set in which the variable was contained. Variables with importance values ≥ 0.8 are considered strongly supported, while those ≥ 0.5 are interpreted as having moderate support, but were considered only, if the respective confidence intervals did not contain zero (Burnham et al., 2011; Symonds & Moussalli, 2011).

In view of the used model-averaging approach, we decided to report effect sizes and their 95% confidence intervals (CIs) for all analyses instead of p-values. Thereby, the interpretation will be based on the magnitude and precision of the biological effects rather than on their significance level (Symonds & Moussalli, 2011).

### 3.2 Modelling the effects of fire history on relevant vegetation structure and abiotic conditions

For our second objective, we examined the influence of the fire-related variables (fire frequency, time since the last fire, maximum fire severity, burn status (burnt vs. unburnt)) on those relevant vegetation structures that were identified in our first objective. We fitted two GLMMs using two datasets for each dependent variable with the “glmmTMB” package. The first dataset (Model A) included all transect partitions across all sites and was used to examine the effects of the burn status (burnt/unburnt) and the number of fires (0-6) on the vegetation parameters and abiotic condition. The second dataset (Model B) included only all burnt partitions and was used to evaluate the impact of the number of years since the last fire and the maximum fire severity on vegetation parameters. Preliminary analyses revealed no significant difference between burnt and transition zones regarding the vegetation parameters (Table S8). Therefore, we combined the transition and burnt transect zones into a single “burnt” category. As our response variables were normally distributed, we selected the Gaussian family for modeling. Both site and month were included as random effect. Results are reported using p-values, with statistical significance determined at p < 0.05.

## 4 RESULTS

We retained three PCs with Eigenvalues ≥ 1 in the PCA that together explained 68.45% of the total variance (Table S1). PC1 (29.67% explained variance) reflects an overstory complexity and closure, with strong positive loadings for tree density (0.354), counts of medium trees (0.307), tall trees (0.461), and very tall trees (0.405), as well as a high cover of the medium (0.463) and high stratum (0.373). PC2 (23.87% explained variance) describes rather the openness of the two lower strata, with negative loadings for small trees (−0.539), medium trees (−0.502), and for small-strata cover (−0.485). Finally, high PC3 values (14.92% explained variance) correspond to habitats with high forest cover at > 10m (0.477) and high counts of very tall trees (0.510) together with a high cover of the understory (0.467) but not of the medium stratum (−0.400) (Table S1). These three PCs were subsequently used as structural predictors in the species-specific GLMMs.

### 4.1 Influence of vegetation structure and abiotic factors on the presence of large-sized lemurs

The analysis identified humidity and tree species richness as influential predictors for the presence of *Eulemur fulvus* (Estimate_humidity_ = 0.07 ± 0.03 SE, 95% CI = [0.002, 0.148], RI = 1.00; Estimate_tree_sp_richness_ = 0.16 ± 0.08 SE, 95% CI = [0.0005, 0.3308], RI = 0.84, Figure 3A). In other words, the probability of *E. fulvus* presence increased with higher humidity levels and higher tree species richness (Figure 3B). In contrast, the vegetation structure PCs (PC1 - PC3) had low importance and confidence intervals overlapped zero (Figure 2, Table S3), while maximum temperature was not even included in the top models (Table S3).

**FIGURE 2.**
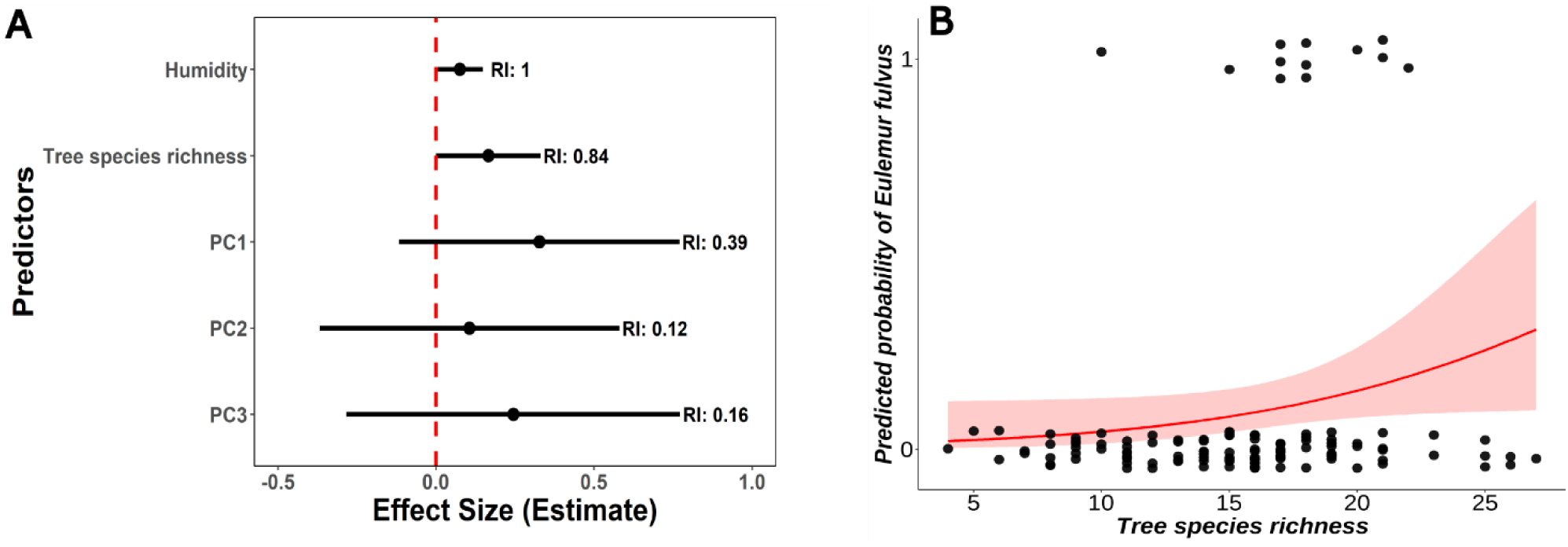
(A) Averaged parameter estimates for predictors of Eulemur fulvus presence that were included in the subset of best models. Points represent effect size estimates on the logit scale, with horizontal whiskers showing 95% confidence intervals. Important variables are those that do not cross the dashed line with their confidence intervals and have an RI ≥ 0.8. Relative Importance (RI) values, based on the sum of Akaike weights across best models, are displayed for each predictor. **(B)** Relationship between tree species richness and the predicted probability of Eulemur fulvus occurrence. Model-generated individual data points are displayed as jitter and fitted probability line (± 95% CI) illustrate a positive association between increasing tree species diversity and the likelihood of E. fulvus presence within surveyed habitats.

**FIGURE 3.**
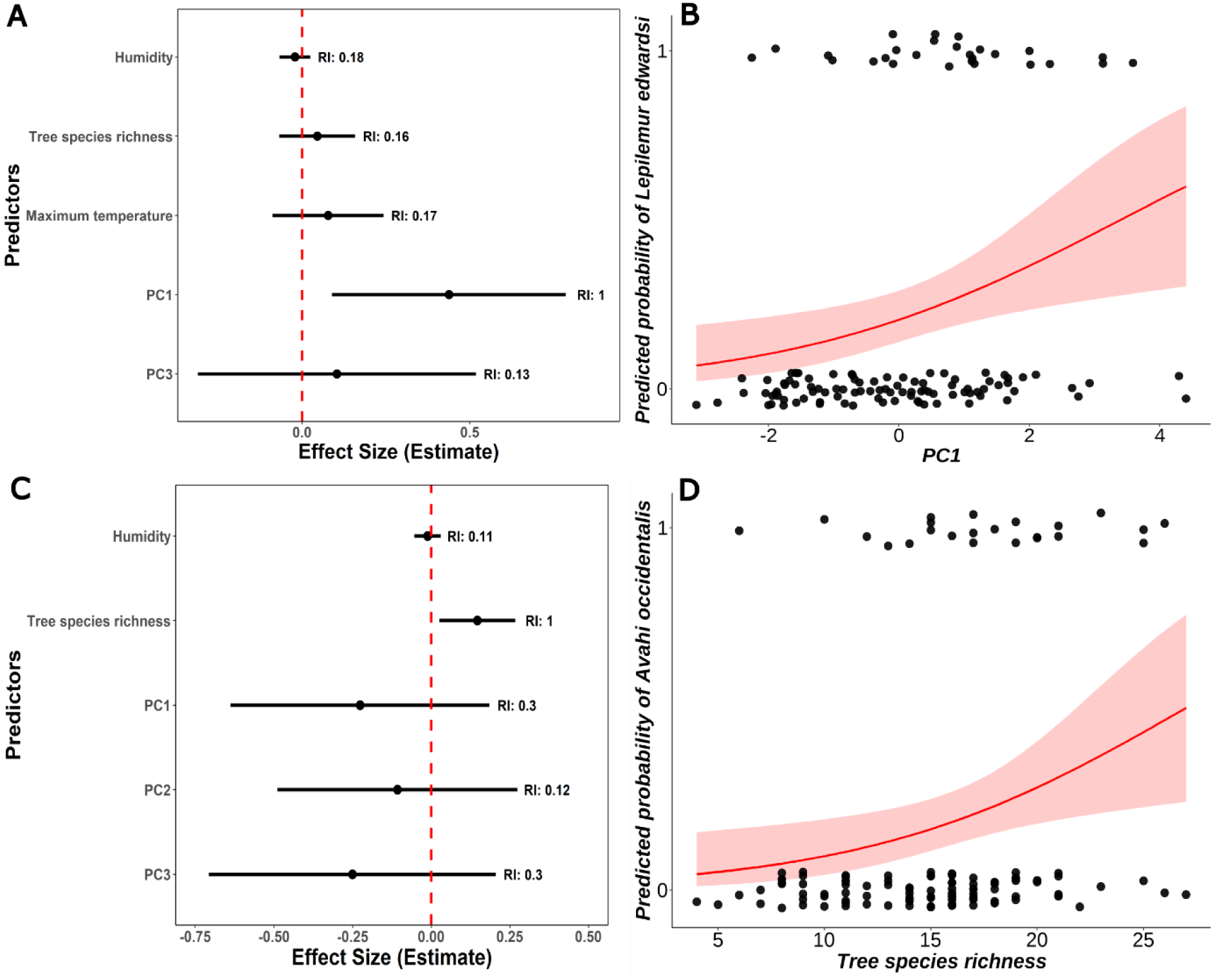
Averaged parameter estimates for predictors for the presence of the two medium-sized lemurs Lepilemur edwardsi **(A)** and Avahi occidentalis **(C).** Effect sizes are displayed on the logit scale for each parameter that was included in the averaged best model, with horizontal whiskers representing 95% confidence intervals. Important variables are those whose confidence intervals do not cross the dashed line and have an RI ≥ 0.8. Relative Importance (RI) values, based on the sum of Akaike weights across best models, are displayed for each predictor. **(B)** Relationship between PC1 and the predicted probability of Lepilemur edwardsi occurrence. Model-generated individual data points are displayed as jitter and fitted probability line (± 95% CI) illustrate a positive association between increasing structural complexity and tall forests and L. edwardsi presence within surveyed habitats. **(D)** Relationship between tree species richness and the predicted probability of Avahi occidentalis occurrence. Model-generated individual data points are displayed as jitter and fitted probability line (± 95% CI) illustrate a positive association between increasing tree diversity and the likelihood of A. occidentalis presence within surveyed habitats.

### 4.2 Influence of vegetation structure and abiotic conditions on the presence of medium-sized lemurs

Forest structural complexity (PC1) was the only important predictor for the presence of *L. edwardsi*, showing a positive relationship with species presence (Estimate_PC1_ = 0.44 ± 0.18 SE, 95% CI = [0.09, 0.79], RI = 1.00, Figure 3A). Since PC1 signaled overstory complexity and closure, *L*. *edwardsi* presence was associated with structurally complex and tall forests (Figure 3B). The other variables (humidity, maximum temperature, tree species richness, PC3) had low importance and confidence intervals overlapped zero, or were not even included in the best model set (PC2, Table S4).

Tree species richness was the only important predictor for the presence of *A. occidentalis*, positively affecting its occurrence probability (Estimate_tree_sp_richness_ = 0.15 ± 0.06 SE, 95% CI = [0.03, 0.27], RI = 1.00, Figures 3C, 3D). Other predictors (PC1, PC3, humidity) showed low relative importance, while maximum temperature was not even included in the best model set (Table S4).

### 4.3 Influence of vegetation structure and abiotic conditions on the presence of small lemurs

Tree species richness had an important positive effect on the probability of occurrence of *Cheirogaleus medius* along the transects (Estimate_tree_sp_richness_ = 0.14 ± 0.07 SE, 95% CI = [0.002, 0.287], RI = 0.83, Figure 4A, 4B). Other predictors, such as PC2, PC3, humidity and maximum temperature were retained in the averaged best model but were not important predictors, while PC1 was not even included in the best averaged model (Table S5). Humidity and PC3 emerged as the strongest predictors of presence for *Microcebus ravelobensis* (Estimate_humidity_ = 0.04 ± 0.01 SE, 95% CI = [0.002, 0.079], RI = 1.00; Estimate_PC3_ = -0.57 ± 0.19 SE, 95% CI = [-0.966, -0.176], RI = 1.00, Figure 4C). Higher occurrence probabilities for this species were therefore associated with more humid forest and with habitats characterized by a dense middle stratum (Figure 4D). The other predictors, namely PC1, PC2, tree species richness, and maximum temperature were not influential (Table S5). In the case of *Microcebus murinus*, no vegetation or abiotic predictor impacted its occurrence probability. All parameters besides PC3 were included in the best averaged model, but their effects were only weak (Figure 4E, Table S5).

**FIGURE 4.**
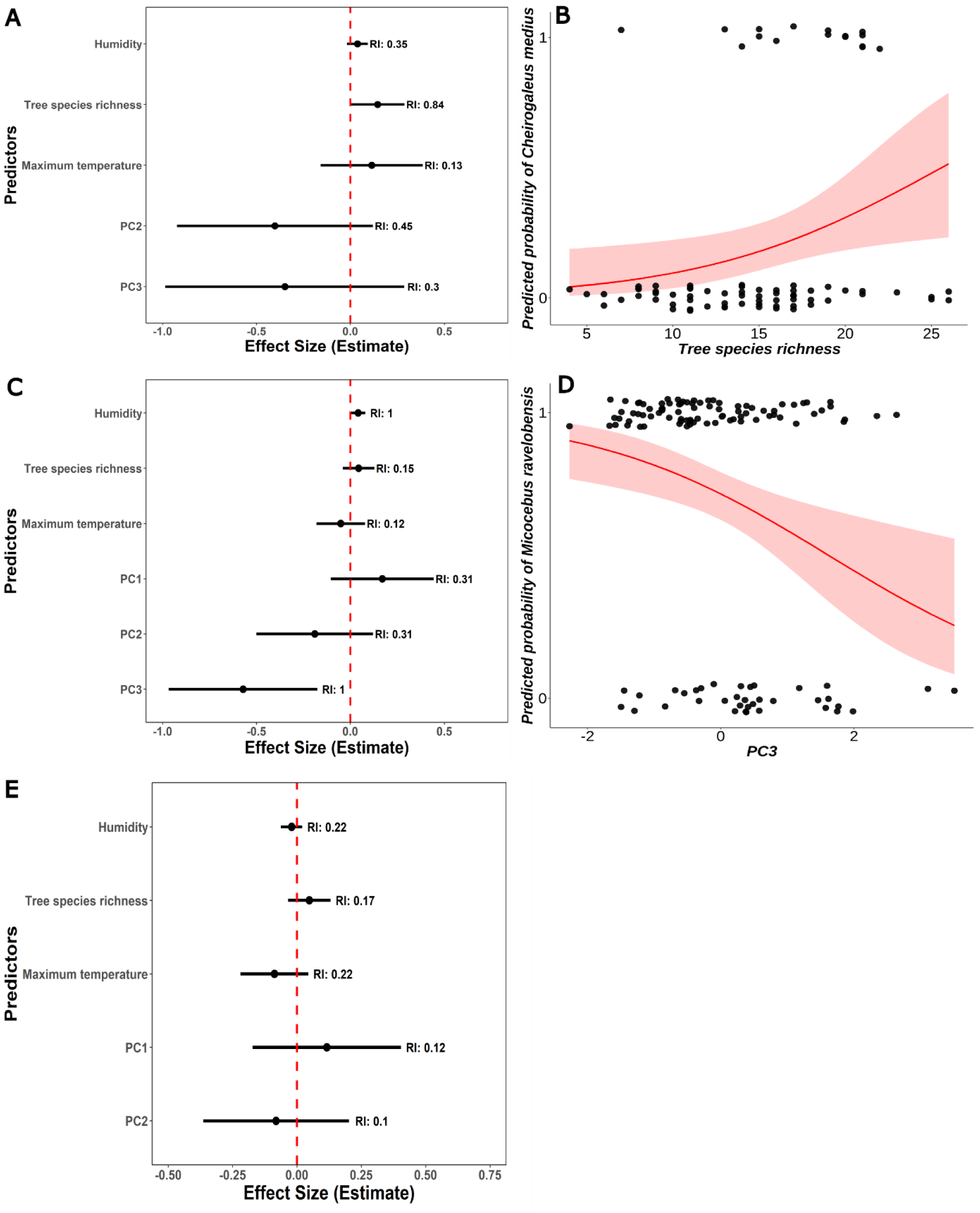
Averaged parameter estimates for predictors for the presence of the small-sized lemurs Cheirogaleus medius **(A)**, Microcebus ravelobensis **(C)**, and Microcebus murinus **(E)**. Effect sizes are displayed on the logit scale for each parameter that was included in the averaged best model, with horizontal whiskers representing 95% confidence intervals. Important variables are those whose confidence intervals do not cross the dashed line and have an RI ≥ 0.8. Relative Importance (RI) values, based on the sum of Akaike weights across best models, are displayed for each predictor. **(B)** Relationship between tree species richness and the predicted probability of Cheirogaleus medius occurrence. Model-generated individual data points are displayed as jitter and fitted probability line (± 95% CI) illustrate a positive association between increasing tree species diversity and C. medius presence within surveyed habitats. **(D)** Relationship between PC3 and the predicted probability of Microcebus ravelobensis occurrence. Model-generated individual data points are displayed as jitter and fitted probability line (± 95% CI) illustrate the relationship between dense middle stratum forest and the likelihood of M. ravelobensis presence within surveyed habitats.

### 4.4 Influence of vegetation structure and abiotic conditions on lemur species richness

Tree species richness emerged as the most important predictor of lemur species richness and had a positive and significant effect (Estimate _tree_sp_richness_ = 0.04 ± 0.01 SE, 95% CI = [0.015, 0.070], RI = 1.00, Figures 5A, 5B). The other predictors, i.e., PC1, PC2, PC3, humidity and maximum temperature, exhibited lower relative importance and confidence intervals overlapping zero, indicating limited explanatory power for lemur species richness (Table S6).

**FIGURE 5.**
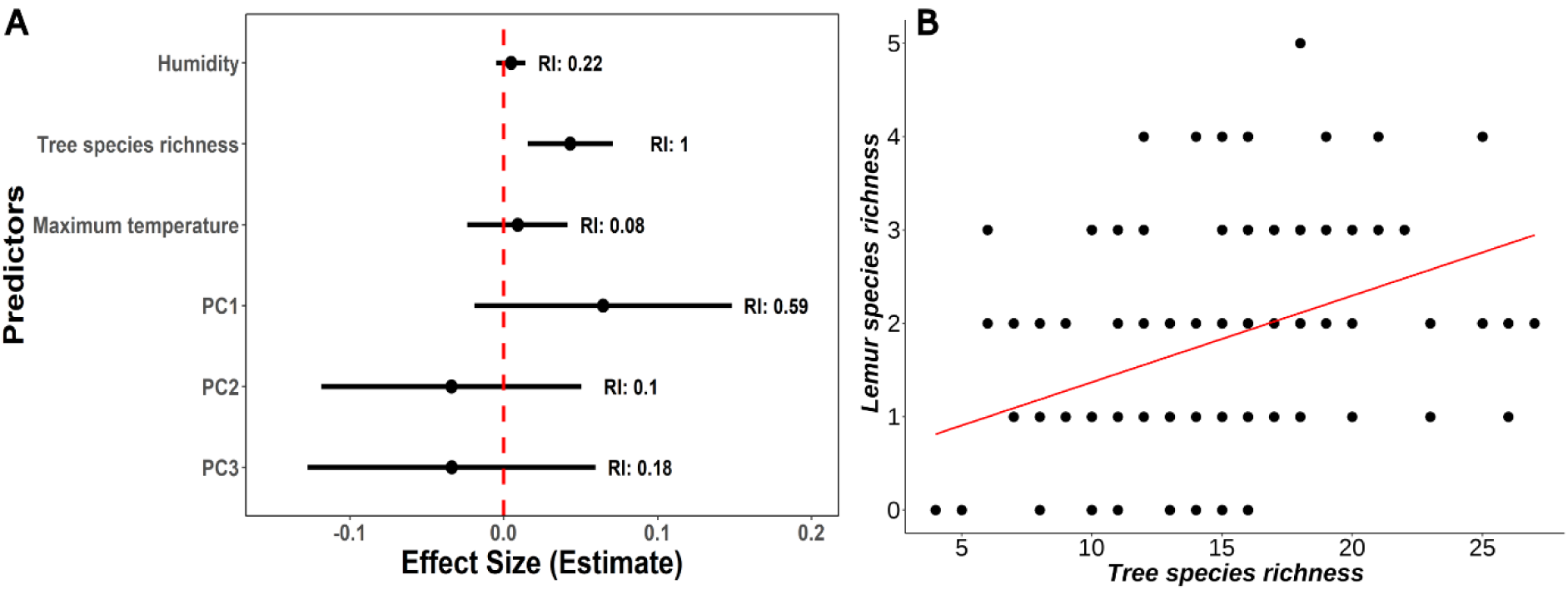
(A) Model-averaged parameter estimates for predictors of lemur species richness. Points represent effect size estimates on the logit scale, with horizontal whiskers indicating 95% confidence intervals. Relative Importance (RI) values, calculated from the sum of Akaike weights across relevant candidate models, are displayed next to each predictor. **(B)** Relationship between tree species richness and the lemur species richness occurrence. Model-generated individual data are displayed (points) and a regression line (± 95% CI) illustrate a positive association between increasing tree species diversity and lemur species richness.

### 4.5 Impact of fire history on relevant vegetation parameters and abiotic factors

#### 4.5.1 Forest humidity and tree species richness

Forest humidity was not strongly influenced by the fire history across all sites. None of the variables showed a detectable association with humidity (Table S8). These results indicate that forest humidity did not differ systematically between burnt and unburned plots and did not clearly respond neither to fire frequency, the time since last fire, nor to the maximum fire severity.

In contrast, tree species richness was affected by the fire history. Specifically, it was higher in unburnt than in burnt plots (Estimate_unburnt_ = 5.05 ± 1.16 SE, 95% CI = [2.77, 7.32], p < 0.0001, Figure 6A), and maximum fire severity had a strong negative effect on tree species richness (Estimate_max_severity_ = -33.1 ± 8.25, 95% CI = [-49.2, -16.9], p < 0.0001, Figure 6B). In contrast, neither the number of fires nor the time since the last fire showed a detectable effect (Table S7).

**FIGURE 6.**
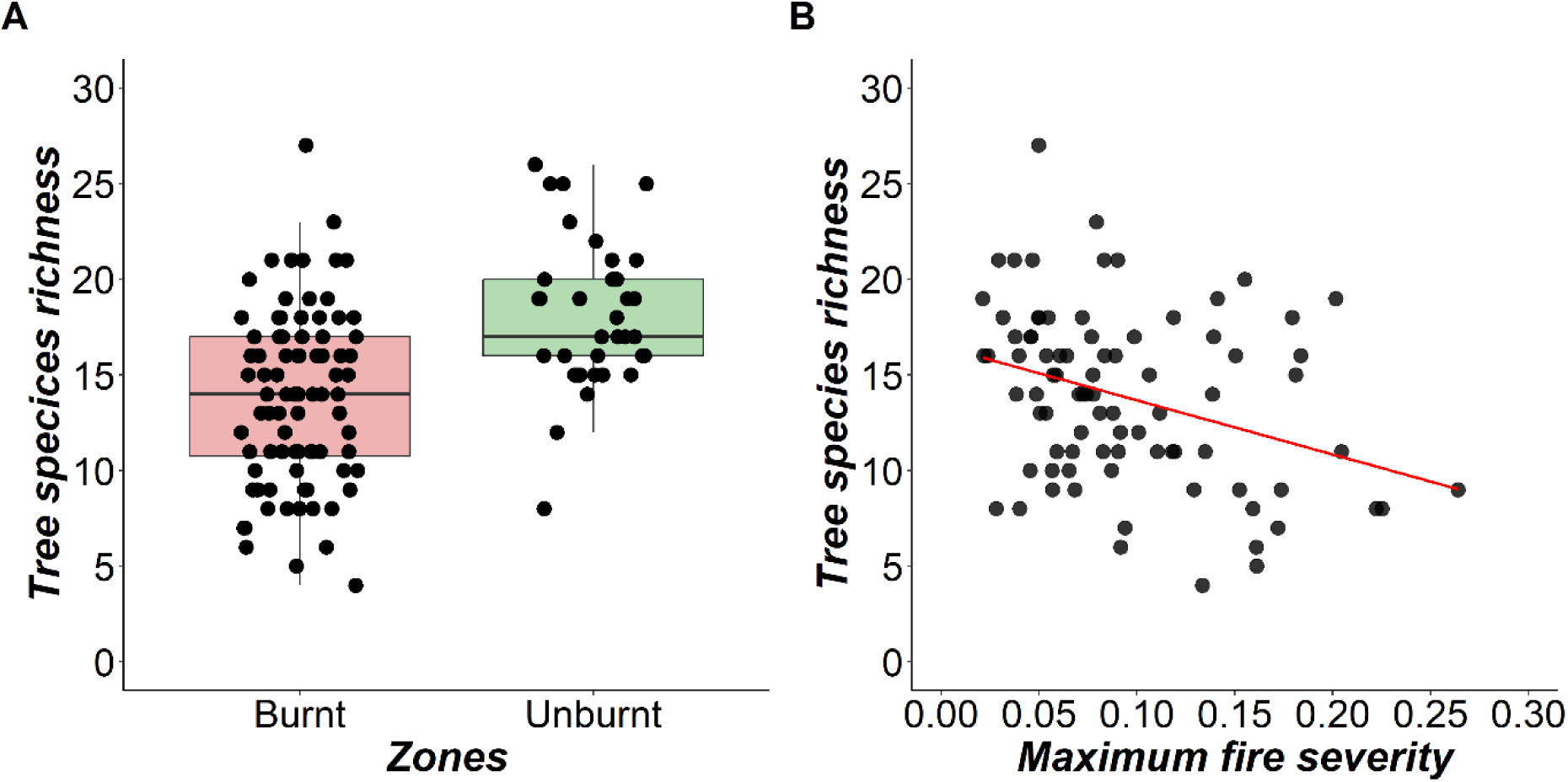
Relationships between tree species richness and **(A)** different fire zones (unburnt vs. burnt) and **(B)** maximum fire severity. The box plot on the left shows the medians and interquartile ranges with individual values being added as jitter. The scatter plot on the right shows individual data points as black dots together with a fitted regression line.

#### 4.5.2 Complexity, density and cover of the vegetation (PC1, PC3)

We modeled the two structural vegetation parameters (PC1, PC3) to assess fire effects on the relevant forest architecture in lemur habitats. PC1 captured overstory complexity and closure and was affected by two fire variables. Unburnt sites had higher PC1 scores (Estimate_unburnt_ = 1.11 ± 0.33 SE, 95% CI = [0.46, 1.75], p < 0.001, Figure 7A), i.e., a higher overstory complexity and closure than burnt sites (Table S8). Moreover, time since the last fire was positively related to PC1 (Estimate_last fire_ = 0.05 ± 0.01 SE, 95% CI = [0.0246, 0.0775], p = 0.0001, Figure 7B), while maximum fire severity had a strong negative effect (Estimate_max_severity_ = -10.5 ± 0.62 SE, 95% CI = [-15.6, -5. 39], p < 0.0001, Figure 7C) showing that severe fires greatly reduced canopy complexity.

**FIGURE 7.**
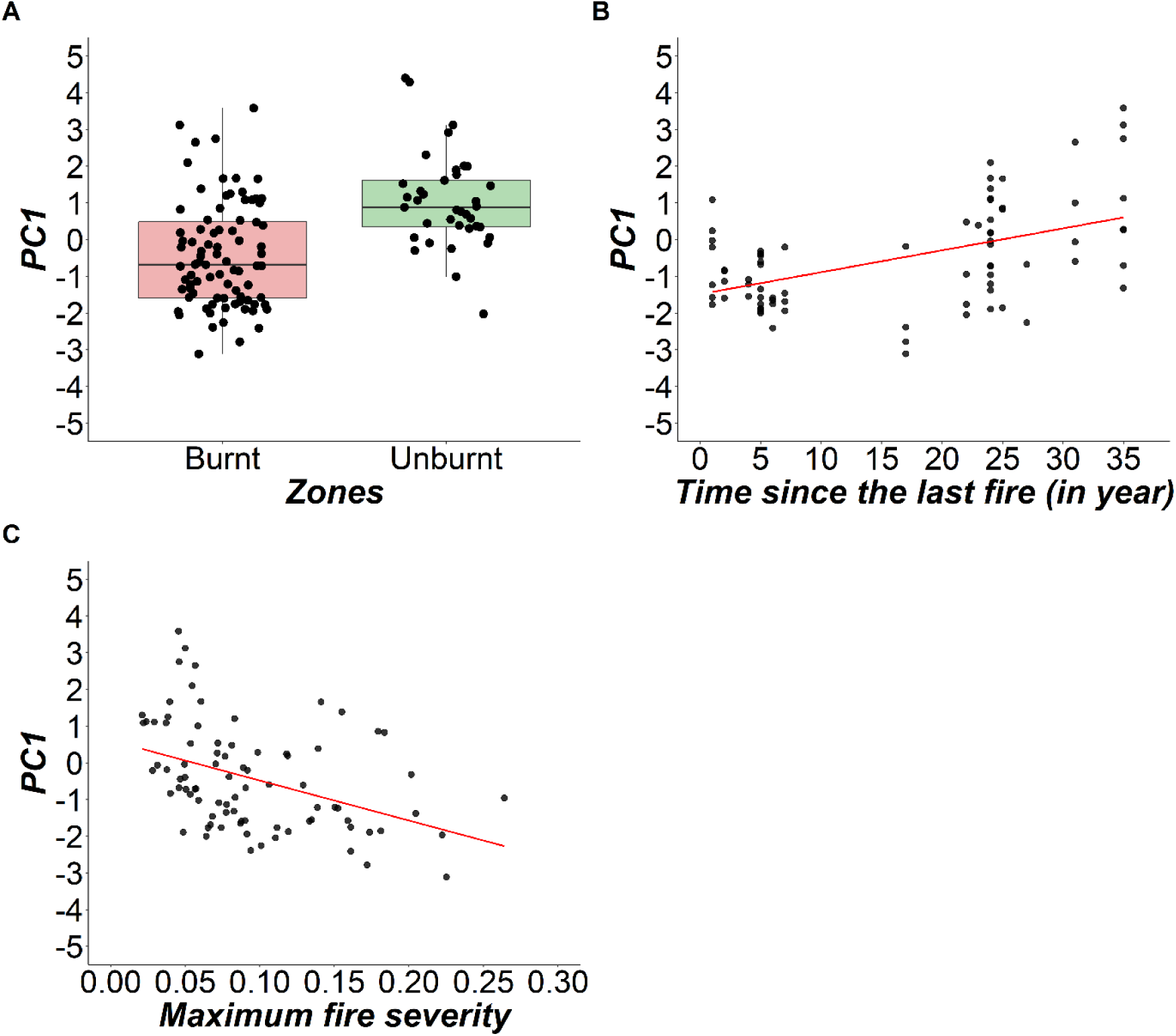
Relationships between PC1 (overstory structural complexity and closure) and influential fire variables. **(A)** Difference in PC1 scores between fire zones (unburnt vs. burnt), illustrated by box plot showing the medians and interquartile ranges; individual datapoints are shown as jitter. **(B, C)** Relationship between PC1 scores and the time since the last fire **(B)** and maximum fire severity (**C)**. Individual data points are shown together with fitted regression line.

Neither burnt status (unburnt/burnt), fire frequency, nor maximum fire severity showed an impact on PC3 (Table S7). Only the time since the last fire showed a small positive influence on PC3, i.e., the density and cover of very tall and small trees (Estimate_last fire_ = 0.02 ± 0.01 SE, 95% CI = [0.002, 0.047], p = 0.03, Figure 8).

**FIGURE 8.**
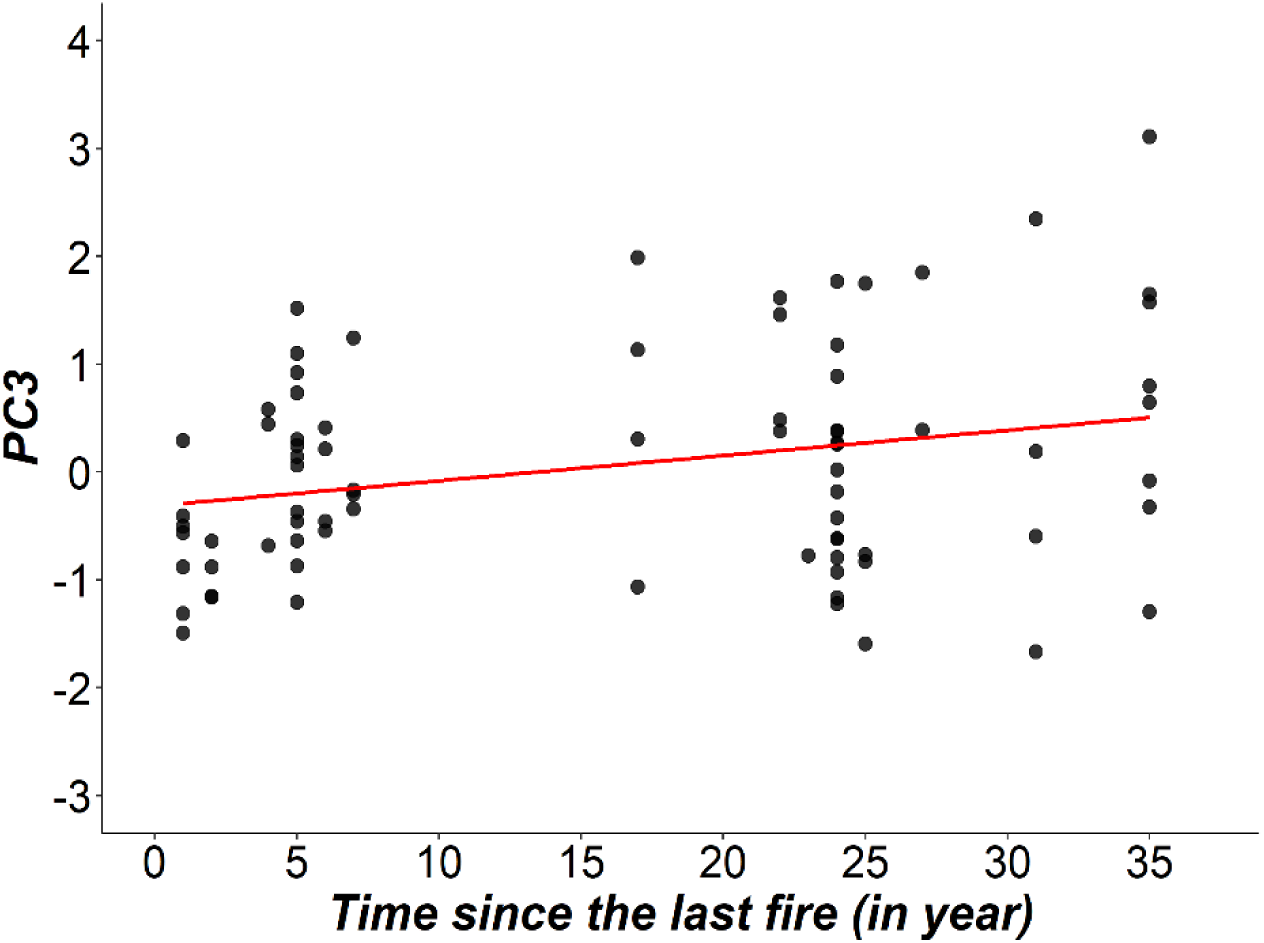
Relationship between PC3 and the time since the last fire. Individual datapoints are shown in black dots together with a fitted regression line.

## 5 DISCUSSION

### 5.1 Impact of fire-modulated vegetation structure and abiotic factors on the large *Eulemur fulvus*

Across Ankarafantsika National Park (ANP), *Eulemur fulvus* exhibited a clear preference for humid forests with high tree species richness. While relative humidity remained consistent across sites with different fire history, tree species diversity peaked in unburnt areas and declined with increasing fire severity. These patterns align with those reported by Rabemananjara et al. (2025), who found *E. fulvus* mainly in unburnt zones or in forests with more than two decades of post-fire recovery. Under such conditions, humid valleys may function as partial fire refugia, where riparian vegetation and higher soil moisture act as natural firebreaks and help preserve essential resources.

As a mainly frugivorous lemur species (Mittermeier et al., 2023), the preference for forests with high tree species richness suggests that *E. fulvus* presence may be constrained by the presence of a diverse spectrum of food species that may provide some fruits even during the lean dry season (Sato, 2013). Intense fires appear to reduce the tree species diversity and thereby negatively influence the occurrence of *E. fulvus*, suggesting that declines in key food resources may limit the species’ ability to persist in burned landscapes. *E. fulvus* may be forced to respond with “habitat shifting”, a strategy that was already observed in response to temporary food scarcity (Sato, 2013). During “habitat shifting”, a species temporarily leaves the established home range to exploit distant habitats with higher food availability (Sato, 2013). Such a strategy is certainly facilitated by the lack of territoriality in *E. fulvus* (Johnson, 2007), although it would likely lead to permanent home range shifts and potential crowding phenomena in case of the rather slow post-fire habitat dynamics (Rabemananjara et al., 2025).

Although limited sampling may have masked some finer-scale associations between vegetation structure and *E. fulvus* occurrence, the species’ association with humid, floristically rich forests suggests that the access to suitable food resources may be a stronger ecological constraint for *E. fulvus* than structural habitat characteristics which were shown to be modified and simplified by fires. However, future studies will be needed to investigate the relationships between food availability, fire history and the presence and abundance of *E. fulvus* in more detail.

### 5.2 Impact of fire-modulated vegetation structure and abiotic factors on medium-sized lemurs

The presence of *Lepilemur edwardsi* in the dry deciduous forest of ANP strongly depended on the availability of vertical forest structures and mature canopy elements (higher values of PC1), but not on tree species richness. With the exception of three sites that had experienced fire 1–5 years earlier, this species was restricted to unburnt areas or to older fire zones where the upper strata had regenerated (Rabemananjara et al., 2025). Our present study further demonstrates that fire-induced loss of large trees (Figure 7C) and canopy simplification reduces the probability of occurrence of *L. edwardsi*. This pattern is consistent with the species’ reliance on suitable substrates for its characteristic clinging-and-leaping locomotion (Warren, 1997) and on secure shelters provided by large overstory trees (Rasoloharijaona et al., 2003; Warren, 1997). Declines in mature tree density (>7 cm DBH) and canopy cover in burned sections may force animals to use midstory routes that carry higher predation risks and energetic costs (Warren & Crompton, 1997). Importantly, Rabemananjara et al. (2025) found that *L. edwardsi* was absent from forests that had experienced intense and more than three fires, indicating a threshold beyond which habitat recovery becomes unlikely or severely delayed. These findings mirror results from other dry forest systems, where high-intensity fires eliminated large trees and destroyed upper forest strata (Smit et al., 2010). Taken together and in contrast to *E. fulvus*, our results suggest that access to appropriate substrates for locomotion and possibly for resting is of greater ecological relevance for this nocturnal folivore than floristic parameters. Future studies should investigate how fire history shapes substrate use and shelter ecology in *L. edwardsi*.

The occurrence of *Avahi occidentalis* was positively related to a higher tree species richness, demonstrating the role of floristic diversity for this specialized folivore (Thalmann, 2001). Fires and fire intensity reduced tree species richness and fires also reduced mature tree density (PC1) which may both have synergistically limited the nutritional options for *A. occidentalis* who depend on diverse diets to cope with toxic plant secondary compounds (Thalmann, 2001). Fire-driven floristic simplification often favors fewer fire-tolerant tree species while reducing the abundance of preferred food plants (Bargali et al., 2022). Our study showed that the fire-related floristic simplification in the Ankarafantsika National Park does severely impact *A. occidentalis* and cannot be compensated by flexible dietary shifts of this species. In this respect, the response of *A. occidentalis* to changes in tree species richness resembles that of *E. fulvus.* It is not yet clear which of the fire-sensitive tree species are keystone resources and drive the strong floristic response of these two lemur species, but this question should be addressed in future work. Rabemananjara et al. (2025) showed that *A. occidentalis* and also *E. fulvus* are more regularly observed again after >23 years post fire, suggesting that tree species richness can recover if no additional fire impacts the forest in the meantime. Such a slow post-fire regeneration has also been observed in other woody assemblages (Allerton et al., 2025).

Overall, the results for *L. edwardsi* and *A. occidentalis* support hypothesis H1 and H2 in a nuanced way. H1 predicted that lemur presence increases with greater structural complexity, canopy connectivity, and floristic diversity, while H2 predicted that these parameters are compromised by fires. While the presence of one medium-sized folivore (*L. edwardsi*) relied mainly on mature, multilayered forests with a well-covered canopy, underscoring the importance of intact vertical structure for their persistence in fire-modulated landscapes, the presence of the other folivore (*A. occidentalis)* depended mainly on a high tree species diversity, i.e., on floristic diversity. This diversity was considerably impacted by past fires, likely constraining the dietary spectrum of available food plants. Finally, neither species was strongly impacted by humidity or temperature variations in the forest. This negligeable role of abiotic factors may be best explained by the long-term territoriality of both species that is maintained despite substantial seasonal variations in abiotic conditions (Ramanankirahina et al., 2012; Rasoloharijaona et al., 2003). It is therefore suggested that changes in temperature or humidity rather lead to changes in microhabitat use, resource use or other adaptive behavioral changes of these two species, but not directly to changes in territory occupancy and therefore presence or absence in a given site.

### 5.3 Impact of fire-modulated vegetation structure and abiotic factors on small lemurs

The probability of occurrence of *Cheirogaleus medius* strongly depended on tree species richness and increased with increasing floristic diversity, which in turn was shown to decline in response to fires and increasing fire severity. Conversely, the other ecological parameters did not show important effects on the presence of *C. medius*. This result resembles closely the findings for *A. occidentalis*, despite their contrasting diet, which is largely omnivorous in the case of *C. medius (*Fietz & Ganzhorn, 1999). However, floristically diverse forests likely provide reliable, though seasonally variable, food resources (e.g., fruits, flowers, seeds, gum, leaves) even during the lean season of the year (Garcia et al., 2014). The availability of a broad spectrum of tree species and food items may be particularly relevant for this lemur species, as it is an obligate hibernator that spends more than five months of the dry season inactive before the beginning of the mating season (Fietz & Ganzhorn, 1999; Lahann, 2007; Souza-Alves et al., 2021). Due to a massive loss in body mass during hibernation (Dausmann et al., 2004), it can be expected that energy requirements will be high after arousal, and that immediate access to energy-rich food resources may have important fitness consequences (Blanco et al., 2021). Under these conditions, relying on and choosing territories that contain a high variety of tree species may provide a reliable “resource insurance”, allowing even to buffer year-to-year uncertainties and variations in food plant phenological dynamics. Our previous work did not find strong impacts of fire variables on the abundance of *C. medius*, although a trend was detected for increasing abundance with increasing fire age (Rabemananjara et al., 2025). However, floristic diversity was not quantified in that study, and the feeding ecology of *C. medius* in this region is still poorly documented. Therefore, further research is needed to clarify whether the presence of *C. medius* in forest sites is primarily driven by the distribution of certain keystone food resources or potentially also by the availability of certain cavity-bearing tree species that facilitate well insulated and protected nesting, torpor and hibernation of small family groups that typically rest and hibernate together during five months of the year (Lahann, 2007; Müller, 1998). Although *C. medius* was observed also in a few burnt and degraded habitats (Hending, 2021; Rabemananjara et al., 2025; Steffens et al., 2020), its presence in these disturbed habitats may represent transient use rather than long-term persistence and may go along with reduced population densities and lower reproductive rates in unfavorable environments (Fietz et al., 2000; Lehman et al., 2006; Schäffler et al., 2015).

The presence of *Microcebus ravelobensis* reflected an affinity to humid microclimates and the importance of a well-covered middle stratum (5-10m) for this species (negative effect of PC3). The importance of mesic conditions for this species was already pointed out in earlier work across the ANP (Rakotondravony & Radespiel, 2009) and largely corresponds to topographically driven microhabitat preferences of this species (Rabemananjara et al., 2025, Steffens et al., 2022). An affinity to a well-developed middle stratum may relate to different aspects of its biology. First, a well-covered middle stratum may contain a large number of suitable sleeping sites for *M. ravelobensis*, as it is known that they use a wide variety of shelter types, among which also dense tangles of lianas and even self-built leaf nests that can be found at these heights, even in unburnt forest (Radespiel et al., 2003, Thorén et al. 2010). Second, a well-covered middle stratum may provide valuable protection from predators for small species like mouse lemurs (Torre et al., 2023). Our study also showed that this structural parameter (PC3) was influenced by the time since the last fire. Specifically, fires led to a higher-cover middle stratum in more recently burned forests which was reduced in favor of the small and large stratum with increasing age of the fire. Such a well-covered middle stratum in recent fire zones may explain why the two smallest lemurs were equally likely found in all sites, whether they were burnt or unburnt (Rabemananjara et al., 2025). This resilience towards fires is consistent with the adaptive habitat flexibility reported in small-bodied lemurs inhabiting disturbed habitats (Hending, 2021). To what extend this habitat flexibility is the result of flexible resource use and dietary shifts in these omnivorous solitary foragers, has to be clarified in future studies.

*Microcebus murinus* also exhibited a strong fire-related ecological resilience, as its probability of occurrence did not vary systematically in response to changes in vegetation structure, tree species diversity, or abiotic conditions. Its persistence across fire-modulated habitats reflects generalist foraging habits and a flexible use of fine-branch microhabitats that persist even in regenerating forests (Hending, 2021). Structural changes and temporary openings in the vegetation may not challenge this species, which readily exploits secondary growth and even small forest fragments for shelter and feeding (Radespiel et al., 2006). Its wide dietary breadth, ranging from arthropods to insect secretions, fruits and gum, further enhances survival in modified post-fire environments (Ganzhorn, 2003; Ganzhorn & Schmid, 1998; Radespiel et al., 2006).

The results on both mouse lemur species align with Hypothesis H2, which predicted that the influence of fire-related vegetation changes on lemur presence varies with body size, i.e., small, generalist lemurs are more tolerant of fire-modulated changes than larger taxa. As a consequence, small-bodied lemur species may function as ecological bridge species, sustaining key processes such as seed dispersal (Ramananjato, 2025), pollination, and invertebrate predation during post-fire recovery phases when larger, canopy-dependent species are absent. Their persistence across varying fire histories underscores the adaptive capacity and ecological resilience of small-bodied generalists.

### 5.4 Lemur species richness: the influence of fire-modulated floristic diversity in structuring lemur assemblages

Because several species were encountered only rarely, a complete assessment of the lemur assemblage was not feasible, thereby constraining our capacity to evaluate fire effects at the community level. The scarcity of observations for the two rarest species (*P. coquereli, E. mongoz*) leaves substantial gaps in our understanding of how fire regimes influence their encounter rates and occurrence. Although patterns derived from the better-sampled taxa provide useful ecological context, they cannot reliably describe community-wide responses. *Propithecus coquereli* was observed predominantly in unburned forest or in areas with fires older than two decades (Rabemananjara et al., 2025). *Eulemur mongoz* exhibited a similarly restricted distribution, occurring along transects at only one of 18 sites and being recorded outside transects at a few additional locations, again almost exclusively in unburned habitat (Rabemananjara et al., 2025). Evaluating their fire responses in more detail will require intensive and sustained monitoring efforts designed to improve detection probabilities for rare species.

Despite these limits, the results for other species already illuminated how fire history shapes lemur habitat use and distribution in the dry deciduous forests of northwestern Madagascar. Tree species richness emerged as the strongest predictor of the overall lemur species richness, underscoring the central role of species-rich plant communities in maintaining multi-species primate assemblages (Ganzhorn, 1999; Schüßler et al., 2018). This relationship was driven by similar patterns observed in three lemur species (*E. fulvus, A. occidentalis, C. medius*) that suggest the importance of diverse food and structural resources, as higher tree species richness provides a wider array of fruiting, flowering, and foliar phenologies that can sustain species with dietary constraints and microhabitat preferences throughout seasonal cycles (Ganzhorn et al., 1997). It can be expected that the real number of vulnerable lemurs in this context is probably even higher, since *P. coquereli* and *E. mongoz* were not included in the modeling. Taken together, our data suggest that tree species richness may potentially serve as an integrative proxy for lemur habitat quality (Huang et al., 2018) given the interdependencies between this parameter and several lemur species.

The negative effect of fire severity on tree species richness observed in this study likely contributed indirectly to the decline in lemur species richness found across frequently burnt or recently disturbed forests (Rabemananjara et al., 2025). Repeated or intense fires simplify forest composition, reducing both vertical stratification and chemical diversity among plant species (Pereira et al., 2025). Such simplification constrains the ecological niches available to specialized lemurs, while favoring a few generalist taxa capable of exploiting secondary vegetation or degraded habitats, such as mouse lemurs. Taken together, these findings reinforce that conserving floristically diverse and structurally complex forests is essential for maintaining full lemur assemblages across fire-prone landscapes.

## 6 CONSERVATION IMPLICATIONS

Our findings from Ankarafantsika National Park underscore that fire is not only a driver of vegetation change but also a critical determinant of lemur community composition. By altering tree diversity, canopy complexity, and floristic composition, recurrent and severe fires progressively erode the ecological foundations that sustain lemur populations. Although only one large-bodied lemur species was represented in our study, its consistent restriction to humid, species-rich forest underscores the vulnerability of this size class to fire-driven habitat degradation. In addition, medium-bodied taxa declined in repeatedly burned areas, even though smaller generalists persist across a wider range of fire histories. Importantly, lemur species richness depends on higher tree species diversity, emphasizing that areas already affected by multiple fires, now widespread across ANP, are likely to support fewer lemur species. As large sections of the protected area have burned in recent decades (Rasolozaka et al., 2025), these spatial trends highlight an urgent conservation concern: fire is progressively concentrating lemur diversity into a shrinking set of less-disturbed forest patches, making the maintenance and protection of these remaining habitats and lemur species increasingly critical.

Given the centrality of vegetation structure and floristic diversity for lemur community organization and the prominent threats imposed by fires to the integrity of the dry deciduous forests of western Madagascar and of ANP (Frappier-Brinton & Lehman, 2022; Rasolozaka et al., 2025), an effective fire management is urgently needed for all protected zones in Madagascar’s dry forest ecosystems. Priority actions should include (1) long-term monitoring of vegetation recovery and lemur assemblage dynamics to evaluate their fire responses and spatio-temporal threats, (2) protection of mesic forested valley systems that may serve as retreat areas for some lemur species, (3) maintenance of complex, mature canopy structure of old growth or long-unburnt forest, (4) promotion of passive forest restoration by preventing recurrence of fires, and (5) fire prevention and recovery planning. Finally, integrated conservation strategies that aim to raise awareness for the destructive ecological effects of forest fires will be essential for maintaining both the lemur diversity and ecological resilience in Madagascar’s increasingly fire-prone dry forest landscapes.

## Acknowledgments

We express our sincere gratitude to the Ministère de l’Environnement et du Développement Durable and the Direction Régionale de l’Environnement (DREDD) Boeny Betsiboka for granting permission. We warmly thank Madagascar National Parks for their unwavering administrative and financial support, especially the General Directors, the financial department, and Madame Lalatiana Odile Randriamiharisoa.

We are also grateful to the former and current Directors of Ankarafantsika National Park, Mandimby Heriniaina Andriambololona and Charles Andriamaniry, as well as the Chef de Volet Opération, the logistical staff, Chefs Secteurs, and all park agents for their assistance during our fieldwork. Our appreciation extends to the Mention Anthropologie et Développement Durable of the Faculty of Sciences at the University of Antananarivo for facilitating all research permits.

We thank our park guides, Jhonny Kennedy and Estoria Una, along with the local community (CLP and Cook) for their valuable help during our site visits. We further acknowledge the support of Marlene Böhm, Johnny Randriafenontsoa, and Fenohery Andriantsitohaina during field data collection.

This project was funded by Madagascar National Parks and supported by the Kreditanstalt für Wiederaufbau (KfW) under contract № 20/DG/DOP/DAFRH/CONV/2022. Additional support for part of the fieldwork was generously provided by the Cologne Zoo, Germany.

## REFERENCES

Allerton, T., Van Bloem, S., & Manlay, R. (2025). Novel Fires Shift Biological Legacies Away From Natural Regeneration in Caribbean Tropical Dry Forest. Journal of Vegetation Science. 10.1111/jvs.70030.

Alonso, L., Schulenberg, T., Radilofe, S., & Missa, O. (2002). A Biological Assessment of the Réserve Naturelle Intégrale d. *Ankarafantsika*, Madagascar. RAP Bulletin of Biological Assessment, 23.

Bargali, H., Calderon, L. P. P., Sundriyal, R. C., & Bhatt, D. (2022). Impact of forest fire frequency on floristic diversity in the forests of Uttarakhand, western Himalaya. *Trees*, Forests and People, 9, 100300. 10.1016/j.tfp.2022.100300.

Bartoń, K. (2022). MuMIn: multi-model inference. R package version 1.46. 0.

Blanco, M., Greene, L., Schopler, R., Williams, C., Lynch, D., Browning, J., Welser, K., Simmons, M., Klopfer, P., & Ehmke, E. (2021). On the modulation and maintenance of hibernation in captive dwarf lemurs. Scientific Reports, 11. 10.1038/s41598-021-84727-3.

Bond, W. J., & Keeley, J. E. (2005). Fire as a global “herbivore”: the ecology and evolution of flammable ecosystems. Trends in Ecology & Evolution, 20(7), 387–394. 10.1016/j.tree.2005.04.025.

Brooks, M. E., Kristensen, K., van Benthem, K. J., Magnusson, A., Berg, C. W., Nielsen, A., Skaug, H. J., Maechler, M., & Bolker, B. M. (2017). Modeling zero-inflated count data with glmmTMB. bioRxiv, 132753.

Burnham, K. P., & Anderson, D. R. (2002). Model selection and multimodel inference: A practical information-theoretic approach (2nd ed.). Springer. 10.1007/b97636.

Burnham, K. P., Anderson, D. R., & Huyvaert, K. P. (2011). AIC model selection and multimodel inference in behavioral ecology: some background, observations, and comparisons. Behavioral Ecology and Sociobiology, 65(1), 23–35. 10.1007/s00265-010-1029-6.

Cant, J. (1992). Positional behavior and body size of arboreal primates: a theoretical framework for field studies and an illustration of its application. American Journal of Physical Anthropology, 88 3, 273–283. 10.1002/AJPA.1330880302 .

Cochrane, M. A. (2003). Fire science for rainforests. Nature, 421(6926), 913–919. 10.1038/nature01437.

Cochrane, M. A., & Laurance, W. F. (2002). Fire as a large-scale edge effect in Amazonian forests. Journal of Tropical Ecology, 18(3), 311–325. 10.1017/S0266467402002237 Cudney-Valenzuela, S., Arroyo-Rodríguez, V., Andresen, E., Toledo-Aceves, T., Mora-Ardila, F.,

Andrade-Ponce, G., & Mandujano, S. (2021). Does patch quality drive arboreal mammal assemblages in fragmented rainforests? Perspectives in Ecology and Conservation, 19, 61–68. 10.1016/J.PECON.2020.12.004.

Cudney-Valenzuela, S., Arroyo-Rodríguez, V., Morante-Filho, J., Toledo-Aceves, T., & Andresen, E. (2022). Tropical forest loss impoverishes arboreal mammal assemblages by increasing tree canopy openness. Ecological applications, 33. 10.1002/eap.2744.

Dausmann, K., Glos, J., Ganzhorn, J., & Heldmaier, G. (2004). Physiology: Hibernation in a tropical primate. Nature, 429, 825–826. 10.1038/429825a.

Dormann, C. F., Calabrese, J. M., Guillera-Arroita, G., Matechou, E., Bahn, V., Bartoń, K., Beale, C. M., Ciuti, S., Elith, J., Gerstner, K., Guelat, J., Keil, P., Lahoz-Monfort, J. J., Pollock, L. J., Reineking, B., Roberts, D. R., Schröder, B., Thuiller, W., Warton, D. I.,…Hartig, F. (2018). Model averaging in ecology: a review of Bayesian, information-theoretic, and tactical approaches for predictive inference. Ecological Monographs, 88(4), 485–504. 10.1002/ecm.1309.

Estrada, A., & Garber, P. A. (2022). Principal Drivers and Conservation Solutions to the Impending Primate Extinction Crisis: Introduction to the Special Issue. International Journal of Primatology, 43(1), 1–14. 10.1007/s10764-022-00283-1.

Fahrig, L. (2017). Ecological Responses to Habitat Fragmentation Per Se. Annual Review of Ecology, Evolution, and Systematics, 48(Volume 48, 2017), 1–23. 10.1146/annurev-ecolsys-110316-022612

Fernandes-Carvalho-De-Andrade, D., & Brazil, B. b. C. A. S. S. B. (2021). Persistent fire effect on forest dynamics and species composition of an old-growth tropical forest. Forest Systems, 30. 10.5424/FS/2021303-16791.

Fietz, J., & Ganzhorn, J. U. (1999). Feeding ecology of the hibernating primate Cheirogaleus medius: how does it get so fat? Oecologia, 121(2), 157–164. 10.1007/s004420050917.

Fietz, J., Zischler, H., Schwiegk, C., Tomiuk, J., Dausmann, K. H., & Ganzhorn, J. U. (2000). High rates of extra-pair young in the pair-living fat-tailed dwarf lemur, Cheirogaleus medius. Behavioral Ecology and Sociobiology, 49(1), 8–17. 10.1007/s002650000269.

Flesch, A. D. (2017). Influence of local and landscape factors on distributional dynamics: a species-centred, fitness-based approach. Proceedings of the Royal Society B: Biological Sciences, 284(1858), 20171001. doi:10.1098/rspb.2017.1001.

Frappier-Brinton, T., & Lehman, S. (2022). The burning island: Spatiotemporal patterns of fire occurrence in Madagascar. PLoS one, 17. 10.1371/journal.pone.0263313.

Ganzhorn, J. (1999). Lemurs as Indicators for Assessing Biodiversity in Forest Ecosystems of Madagascar: Why it does not Work. 163–174. 10.1007/978-94-011-4677-7_10.

Ganzhorn, J. U. (2003). Effects of introduced Rattus rattus on endemic small mammals in dry deciduous forest fragments of western Madagascar. Animal Conservation, 6(2), 147–157. 10.1017/S1367943003003196.

Ganzhorn, J. U., Malcomber, S., Andrianantoanina, O., & Goodman, S. M. (1997). Habitat Characteristics and Lemur Species Richness in Madagascar. Biotropica, 29(3), 331–343. 10.1111/j.1744-7429.1997.tb00434.x.

Ganzhorn, J. U., & Schmid, J. (1998). Different Population Dynamics of Microcebus murinus in Primary and Secondary Deciduous Dry Forests of Madagascar. International Journal of Primatology, 19(5), 785–796. 10.1023/A:1020337211827.

Garcia, L., Hobbs, R., Santos, F. M. D., & Rodrigues, R. (2014). Flower and Fruit Availability along a Forest Restoration Gradient. Biotropica, 46. 10.1111/btp.12080.

Gautier, L., Tahinarivony, J., Ranirison, P., Wohlhauser, S., Goodman, S., & Raherilalao, M. (2018). The terrestrial protected areas of Madagascar: their history, description, and biota. In.

Gestich, C. C., Arroyo-Rodríguez, V., Saranholi, B. H., da Cunha, R. G. T., Setz, E. Z. F., & Ribeiro, M. C. (2022). Forest loss and fragmentation can promote the crowding effect in a forest-specialist primate. Landscape Ecology, 37(1), 147–157. 10.1007/s10980-021-01336-1.

González, T., González-Trujillo, J. D., Muñoz, A., & Armenteras, D. (2022). Effects of fire history on animal communities: a systematic review. Ecological Processes, 11. 10.1186/s13717-021-00357-7.

Goodman, S. M., & Benstead, J. P. (2005). Updated estimates of biotic diversity and endemism for Madagascar. Oryx, 39(1), 73–77. 10.1017/S0030605305000128.

Goodman, S. M., Raherilalao, M. J., & Wohlhauser, S. (2021). The Terrestrial Protected Areas of Madagascar: Their History, Description, and Biota, Volume 3: Western and Southwestern Madagascar-Synthesis. Association Vahatra.

Harcourt, A., & Doherty, D. (2005). Species–area relationships of primates in tropical forest fragments: a global analysis. Journal of Applied Ecology, 42, 630–637. 10.1111/J.1365-2664.2005.01037.X.

Harrison, M., Deere, N., Imron, M., Nasir, D., Adul, Asti, H. A., Soler, J. A., Boyd, N., Cheyne, S., Collins, S., D’Arcy, L., Erb, W., Green, H., Healy, W., Hendri, Holly, B., Houlihan, P., Husson, S., Iwan,…Struebig, M. (2024). Impacts of fire and prospects for recovery in a tropical peat forest ecosystem. Proceedings of the National Academy of Sciences of the United States of America, 121. 10.1073/pnas.2307216121.

Hartig, F., & Hartig, M. (2022). Package ‘dharma’. *R Package. Available online:* https://CRAN.R-project.org/package=DHARMa (accessed on 5 September 2022).

Hending, D. (2021). Environmental drivers of Cheirogaleidae population density: Remarkable resilience of Madagascar’s smallest lemurs to habitat degradation. Ecology and Evolution, 11(11), 5874–5891. 10.1002/ece3.7449.

Huang, Y., Chen, Y., Castro-Izaguirre, N., Baruffol, M., Brezzi, M., Lang, A., Li, Y., Härdtle, W., Von Oheimb, G., Yang, X., Liu, X., Pei, K.-Q., Both, S., Yang, B., Eichenberg, D., Assmann, T., Bauhus, J., Behrens, T., Buscot, F.,…Schmid, B. (2018). Impacts of species richness on productivity in a large-scale subtropical forest experiment. science, 362, 80–83. 10.1126/science.aat6405.

Ishii, H. T., Tanabe, S.-i., & Hiura, T. (2004). Exploring the Relationships Among Canopy Structure, Stand Productivity, and Biodiversity of Temperate Forest Ecosystems. Forest Science, 50(3), 342–355.

Johns, A., & Skorupa, J. (1987). Responses of rain-forest primates to habitat disturbance: A review. International Journal of Primatology, 8, 157–191. 10.1007/BF02735162

Johnsingh, A. (1986). Impact of fire on wildlife ecology in two dry deciduous forests in south India. Indian Forester, 112(10), 933–938.

Johnson, S. E. (2007). Evolutionary Divergence in the Brown Lemur Species Complex. In L. Gould & M. L. Sauther (Eds.), Lemurs: Ecology and Adaptation (pp. 187–210). Springer US. 10.1007/978-0-387-34586-4_9.

Joseph, G., Seymour, C., & Rakotoarivelo, A. (2023). Fire incongruities can explain widespread landscape degradation in Madagascar’s forests and grasslands. *PLANTS, PEOPLE*, PLANET. 10.1002/ppp3.10471.

Keeley, J. E. (2009). Fire intensity, fire severity and burn severity: a brief review and suggested usage. International Journal of Wildland Fire, 18(1), 116–126. 10.1071/WF07049.

Keeley, J. E., Bond, W. J., Bradstock, R. A., Pausas, J. G., & Rundel, P. W. (2011). Fire in Mediterranean Ecosystems: Ecology, Evolution and Management. Cambridge University Press. DOI: 10.1017/CBO9781139033091.

Key, C. H., & Benson, N. C. (2006). Landscape assessment (LA). FIREMON: Fire effects monitoring and inventory system, 164, LA–1–55.

Kull, C. A. (2004). Isle of fire: the political ecology of landscape burning in Madagascar. University of Chicago press.

Lahann, P. (2007). Feeding ecology and seed dispersal of sympatric cheirogaleid lemurs (Microcebus murinus, Cheirogaleus medius, Cheirogaleus major) in the littoral rainforest of south-east Madagascar. Journal of Zoology, 271(1), 88–98. 10.1111/j.1469-7998.2006.00222.x.

LaRue, E. A., Hardiman, B. S., Elliott, J. M., & Fei, S. (2019). Structural diversity as a predictor of ecosystem function. Environmental Research Letters, 14(11), 114011. 10.1088/1748-9326/ab49bb.

Le, T. M., & Clarke, B. S. (2022). Model averaging is asymptotically better than model selection for prediction. Journal of Machine Learning Research, 23(33), 1–53.

Lehman, S. M., Rajaonson, A., & Day, S. (2006). Edge Effects on the Density of Cheirogaleus major. International Journal of Primatology, 27(6), 1569–1588. 10.1007/s10764-006-9099-z.

Lenth, R. (2023). emmeans: Estimated Marginal Means, aka Least-Squares Means_. R package version 1.8. 5.

Leys, B. A., Marlon, J. R., Umbanhowar, C., & Vannière, B. (2018). Global fire history of grassland biomes. Ecology and Evolution, 8(17), 8831–8852. 10.1002/ece3.4394.

Lindenmayer, D., Blanchard, W., Blair, D., McBurney, L., & Banks, S. (2017). Relationships between tree size and occupancy by cavity-dependent arboreal marsupials. Forest Ecology and Management, 391, 221–229. 10.1016/J.FORECO.2017.02.014.

Lyon, L., Huff, M., Hooper, R., Telfer, E., Schreiner, D., & Smith, J. (2012). Wildland Fire in Ecosystems: Effects of Fire on Fauna. 10.2737/rmrs-gtr-42-v1.

McLean, K. A., Trainor, A. M., Asner, G. P., Crofoot, M. C., Hopkins, M. E., Campbell, C. J., Martin, R. E., Knapp, D. E., & Jansen, P. A. (2016). Movement patterns of three arboreal primates in a Neotropical moist forest explained by LiDAR-estimated canopy structure. Landscape Ecology, 31(8), 1849–1862. 10.1007/s10980-016-0367-9.

Mittermeier, R. A., Reuter, K. R., Rylands, A. B., Louis Jr., E. E., Ratsimbazafy, J., Rene de Roland, L.-A., Langrand, O., Schwitzer, C., Johnson, S. E., Godfrey, L. R., Blanco, M. B., Borgerson, C., Eppley, T. M., Andriamanana, T., Volampeno, S., Andriantsaralaza, S., Wright, P. C., & Rajaobelina, S. (2023). Lemurs of Madagascar. (5th edition ed., pp. 975): *Re:wild*.

Moser, B., Temperli, C., Schneiter, G., & Wohlgemuth, T. (2010). Potential shift in tree species composition after interaction of fire and drought in the Central Alps. European Journal of Forest Research, 129, 625–633. 10.1007/s10342-010-0363-6.

Müller, A. E. (1998). A preliminary report on the social organisation of Cheirogaleus medius (Cheirogaleidae; Primates) in north-west Madagascar. Folia Primatologica, 69(3), 160–166.

Müller, A. E., & Thalmann, U. (2002). Biology of the fat-tailed dwarf lemur (Cheirogaleus medius E. Geoffroy 1812): new results from the field. Evolutionary Anthropology: Issues, News, and Reviews: Issues, News, and Reviews, 11(S1), 79–82.

Nolan, R., Blackman, C., De Dios, V. R., Choat, B., Medlyn, B., Li, X., Bradstock, R., & Boer, M. (2020). Linking Forest Flammability and Plant Vulnerability to Drought. Forests. 10.3390/f11070779.

Pereira, C. A., Barlow, J., Tabarelli, M., Giles, A., De Melo Ferreira, A. E., & Vieira, I. (2024). Recurrent wildfires alter forest structure and community composition of terra firme Amazonian forests. Environmental Research Letters, 19. 10.1088/1748-9326/ad77e6.

Pereira, M. B., Elias, F., Teixeira, N. D. A., Feldpausch, T. R., Marimon-Junior, B. H., & Marimon, B. S. (2025). Post-fire changes in tree diversity, composition and carbon in seasonal forests in the Southern Amazonia. Forest Ecology and Management, 578, 122447. 10.1016/j.foreco.2024.122447.

Peres, C. A. (1999). General guidelines for standardizing line-transect surveys of tropical forest primates. Neotropical primates, 7(1), 11–16.

Phelps, L. N., Andela, N., Gravey, M., Davis, D. S., Kull, C. A., Douglass, K., & Lehmann, C. E. (2022). Madagascar’s fire regimes challenge global assumptions about landscape degradation. Global Change Biology, 28(23), 6944–6960.

Posit team. (2025). RStudio: Integrated Development Environment for R. In http://www.posit.co/

R Core Team. (2025). R: A Language and Environment for Statistical Computing. In R Foundation for Statistical Computing. https://www.R-project.org/.

Rabemananjara, N. R., Rasolozaka, M. M., Ravolanirina, M. O., Marivola, R., Randriamiarantsoa, S. H., Rakotondravony, R., Razafindraibe, H., Schüßler, D., & Radespiel, U. (2025). Post-fire recolonization of dry deciduous forests by lemurs in northwestern Madagascar. Biodiversity and Conservation. 10.1007/s10531-025-03167-x.

Radespiel, U., Ehresmann, P., & Zimmermann, E. (2003). Species-specific usage of sleeping sites in two sympatric mouse lemur species (Microcebus murinus and M. ravelobensis) in northwestern Madagascar. American Journal of Primatology, 59(4), 139–151. 10.1002/ajp.10071.

Radespiel, U., Reimann, W., Rahelinirina, M., & Zimmermann, E. (2006). Feeding Ecology of Sympatric Mouse Lemur Species in Northwestern Madagascar. International Journal of Primatology, 27(1), 311–321. 10.1007/s10764-005-9005-0.

Rakotondravony, R., & Radespiel, U. (2007). Variation de la distribution de deux espèces de microcebes dans le Parc National Ankarafantsika [SCIENCES ANIMALES]. Lemur News, 12, 31–34.

Rakotondravony, R., & Radespiel, U. (2009). Varying patterns of coexistence of two mouse lemur species (Microcebus ravelobensis and M. murinus) in a heterogeneous landscape. American Journal of Primatology: Official Journal of the American Society of Primatologists, 71(11), 928–938.

Ramananjato, V. (2025). Contribution of small nocturnal lemurs to seed dispersal in Madagascar: A review. Biotropica, 57(1), e13379. 10.1111/btp.13379.

Ramanankirahina, R., Joly, M., & Zimmermann, E. (2012). Seasonal Effects on Sleeping Site Ecology in a Nocturnal Pair-Living Lemur (Avahi occidentalis). International Journal of Primatology, 33(2), 428–439. 10.1007/s10764-012-9587-2.

Rasoloharijaona, S., Rakotosamimanana, B., Randrianambinina, B., & Zimmermann, E. (2003). Pair-specific usage of sleeping sites and their implications for social organization in a nocturnal Malagasy primate, the Milne Edwards’ sportive lemur (Lepilemur edwardsi). American Journal of Physical Anthropology, 122(3), 251–258. 10.1002/ajpa.10281

Rasolozaka, M., Schüßler, D., Randriafenontsoa, J., Andriatsitohaina, F., Rakotomamonjy, P., Rabarison, H., & Radespiel, U. (2025). Reconstructing the alarming fire history of Ankarafantsika National Park in northwestern Madagascar over a 35 year-period. Remote Sensing Applications: Society and Environment, 38, 101521. 10.1016/j.rsase.2025.101521.

Rodríguez-Gómez, G., & Fontúrbel, F. (2020). Regional-scale variation on Dromiciops gliroides occurrence, abundance, and activity patterns along a habitat disturbance gradient. Journal of Mammalogy, 101, 733–741. 10.1093/jmammal/gyaa022.

Sato, H. (2013). Habitat shifting by the common brown lemur (Eulemur fulvus fulvus): a response to food scarcity. Primates, 54(3), 229–235. 10.1007/s10329-013-0353-7.

Schäffler, L., Saborowski, J., & Kappeler, P. M. (2015). Agent-mediated spatial storage effect in heterogeneous habitat stabilizes competitive mouse lemur coexistence in Menabe Central, Western Madagascar. BMC ecology, 15(1), 7. 10.1186/s12898-015-0040-1

Schüßler, D., Radespiel, U., Ratsimbazafy, J., & Mantilla-Contreras, J. (2018). Lemurs in a dying forest: Factors influencing lemur diversity and distribution in forest remnants of north-eastern Madagascar. Biological Conservation. 10.1016/j.biocon.2018.10.008

Schwitzer, C., Mittermeier, R. A., Davies, N., Johnson, S., Ratsimbazafy, J. R., J, Louis Jr., E. E., & Rajaobelina, S. (2013). Lemurs of Madagascar a strategy for their conservation 2013-2016. IUCN SSC Primate Specialist Group, Bristol Conservation and Science Foundation, and Conservation International, UK.

Smit, I. P. J., Asner, G. P., Govender, N., Kennedy-Bowdoin, T., Knapp, D. E., & Jacobson, J. (2010). Effects of fire on woody vegetation structure in African savanna. Ecological applications, 20(7), 1865–1875. 10.1890/09-0929.1.

Souza-Alves, J. P., Chagas, R. R. D., Santana, M. M., Boyle, S. A., & Bezerra, B. M. (2021). Food availability, plant diversity, and vegetation structure drive behavioral and ecological variation in Endangered Coimbra-Filho’s titi monkeys. American Journal of Primatology, 83(3), e23237. 10.1002/ajp.23237.

Steffens, T. S., Mercado Malabet, F., & Lehman, S. M. (2020). Occurrence of lemurs in landscapes and their species-specific scale responses to habitat loss. American Journal of Primatology, 82(4), e23110.

Symonds, M. R. E., & Moussalli, A. (2011). A brief guide to model selection, multimodel inference and model averaging in behavioural ecology using Akaike’s information criterion. Behavioral Ecology and Sociobiology, 65(1), 13–21. 10.1007/s00265-010-1037-6.

Thalmann, U. (2001). Food Resource Characteristics in Two Nocturnal Lemurs with Different Social Behavior: Avahi occidentalis and Lepilemur edwardsi. International Journal of Primatology, 22(2), 287–324. 10.1023/A:1005627732561.

Torre, I., Ribas, A., & Puig-Gironès, R. (2023). Effects of Post-Fire Management on a Mediterranean Small Mammal Community. Fire, 6(1), 34.

Warren, R. D. (1997). Habitat use and support preference of two free-ranging saltatory lemurs (Lepilemur edwardsi and Avahi occidentalis). Journal of Zoology, 241(2), 325–341. 10.1111/j.1469-7998.1997.tb01963.x.

Warren, R. D., & Crompton, R. H. (1997). Locomotor ecology of Lepilemur edwardsi and Avahi occidentalis. American Journal of Physical Anthropology, 104(4), 471–486. 10.1002/(SICI)1096-8644(199712)104:4<471::AID-AJPA4>3.0.CO;2-V.

