## Supplementary material for "Influence of vegetation structure on lemur recolonization of post-fire habitats in northwestern Madagascar": Suplementary information

**Supplementary Figures**

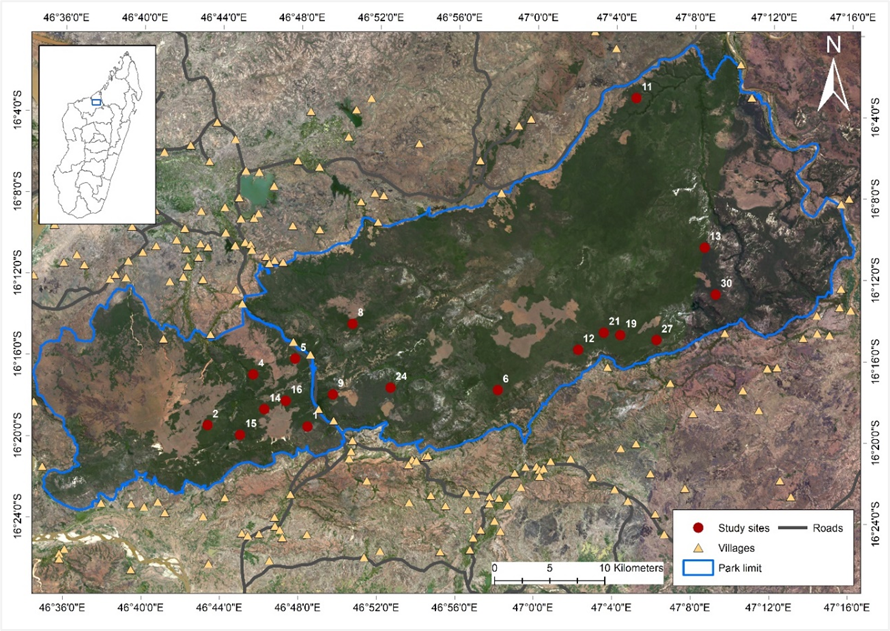

**FIGURE S1** Map of Ankarafantsika National Park located in northwestern Madagascar (inlay map) with the 18 study sites selected during prior remote sensing work (Source: Rabemananjara et al., 2025)

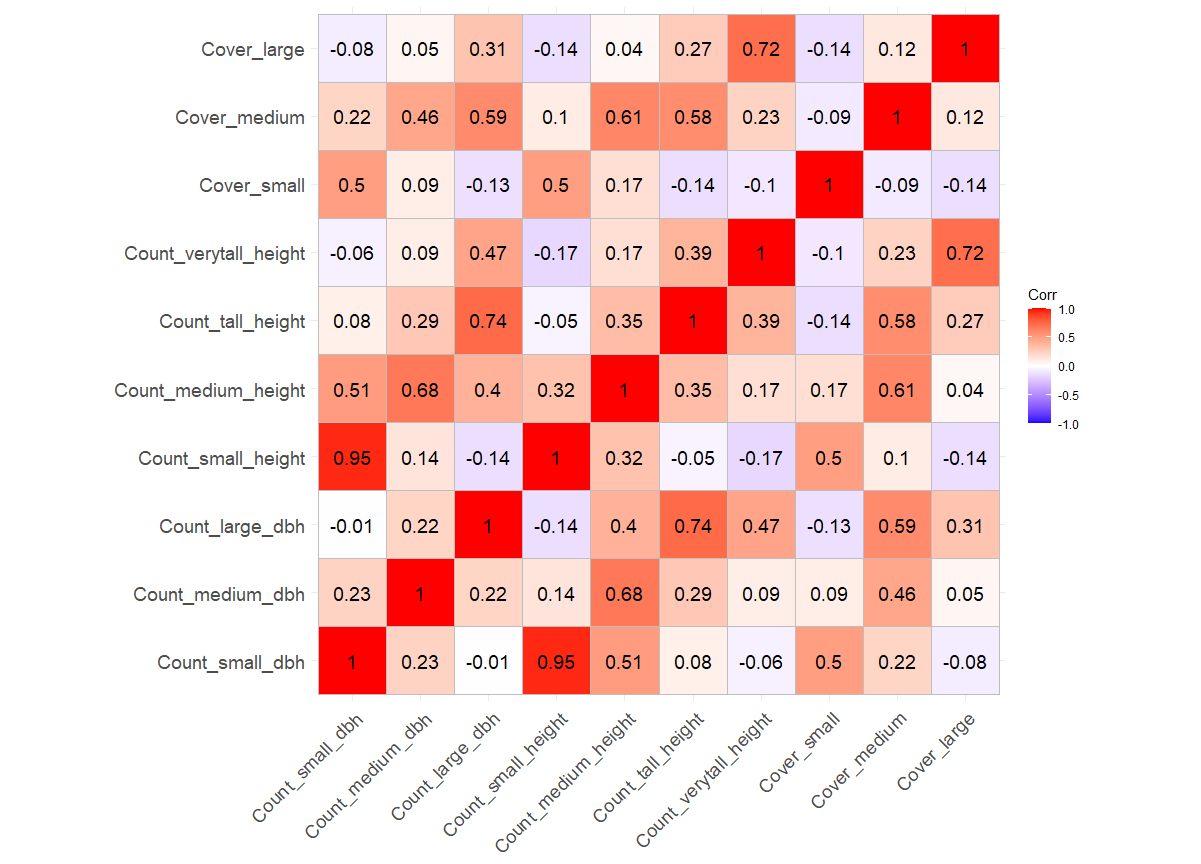

**FIGURE S2** Correlation heatmap of vegetation structural variables (DBH class, Height class, and forest strata cover class). Correlated variables with absolute values ≥ 0.7 were not included for PCA.

**Supplementary Tables**

**TABLE S1** Summary of the principal component analysis (PCA) for vegetation-structure variables. Only components with eigenvalues greater than 1 were retained for subsequent analyses. The table reports the eigenvalues, the proportion of variance explained, and the cumulative proportion of variance explained by each principal component. The first three components together account for 68.5% of the total variance. Loadings of the vegetation-structure variables on each principal component are also presented. Variables with loadings > 0.3 were considered to contribute meaningfully to a component and were highlighted in grey for PC1–PC3, which were included in further analyses. PC1 reflects overstory complexity and canopy closure. PC2 primarily describes openness in the two lower strata, with negative loadings for small trees. PC3 corresponds to habitats characterized by high forest cover above 10 m and high counts of very tall trees, combined with a dense understory but low cover in the medium stratum.

|  | PC1 | PC2 | PC3 | PC4 | PC5 | PC6 | PC7 | PC8 |
| --- | --- | --- | --- | --- | --- | --- | --- | --- |
| Eigenvalue | 1.541066 | 1.382385 | 1.093097 | 0.932705 | 0.782867 | 0.694038 | 0.543952 | 0.514399 |
| Proportion of explained variance | 0.29665 | 0.2387 | 0.14925 | 0.10866 | 0.07656 | 0.06017 | 0.03696 | 0.03305 |
| Cumulative Proportion | 0.29665 | 0.53535 | 0.6846 | 0.79327 | 0.86982 | 0.92999 | 0.96695 | 1 |
| Tree density | 0.354342204 | -0.235898451 | 0.146874254 | -0.708564149 | -0.344122725 | -0.2938651 | 0.142785583 | 0.264562758 |
| Count of small trees | -0.143940239 | -0.539336606 | 0.224224387 | 0.450535037 | 0.143702667 | -0.40612261 | 0.063719592 | 0.495472118 |
| Count of medium trees | 0.30668912 | -0.501985849 | -0.130911211 | -0.171784534 | 0.46922266 | 0.08094985 | -0.568967323 | -0.238448575 |
| Count of tall trees | 0.46094454 | 0.018803492 | -0.212752516 | 0.423291585 | -0.610448691 | 0.06759134 | -0.403710953 | 0.150128054 |
| Count of very tall trees | 0.40451384 | 0.181851619 | 0.510042755 | 0.047472787 | 0.280274922 | 0.54624921 | 0.067445588 | 0.39926197 |
| Cover of small stratum | -0.172569102 | -0.484720044 | 0.46650745 | 0.070433109 | -0.403611367 | 0.3683627 | 0.11356641 | -0.448537417 |
| Cover of medium stratum | 0.463313898 | -0.23650935 | -0.400173058 | 0.200846807 | 0.129916295 | 0.07716795 | 0.686362524 | -0.187080176 |
| Cover of large stratum | 0.372992513 | 0.277784767 | 0.477324533 | 0.196762308 | 0.098879591 | -0.54548335 | -0.017492617 | -0.457736654 |

**TABLE S2** Model selection results showing the best-fitting models for different lemur species and lemur species richness, with associated model parameters and selection criteria. The table includes the species assessed, the component model representing the set of predictor variables, the degrees of freedom (df), the log-likelihood (logLik), the corrected Akaike Information Criterion (AICc) which balances fit and complexity and is corrected for small sample sizes with lower values representing better models, the delta (ΔAICc) which is the difference between a given model and the best-fitting model, and the Akaike weight which reflects the relative likelihood of a model being the best among the candidate models. Null: null model that does not contain any predictor variable. PC: principal component.

| Species | Component model | df | logLik | AICc | Delta (AICc) | Akaike Weight |
| --- | --- | --- | --- | --- | --- | --- |
| *Eulemur fulvus* | Humidity + Tree species richness | 4 | -33.78 | 75.9 | 0.0 | 0.32 |
|  | Humidity + Tree species richness + PC1 | 5 | -33.0 | 76.53 | 0.63 | 0.24 |
|  | Humidity +Tree species richness + PC3 | 5 | -33.36 | 77.25 | 1.34 | 0.16 |
|  | Humidity + PC1 | 4 | -34.49 | 77.33 | 1.42 | 0.16 |
|  | Humidity + Tree species richness + PC2 | 5 | -33.68 | 77.88 | 1.98 | 0.12 |
| *Lepilemur edwardsi* | PC1 | 3 | -59.79 | 125.79 | 0.0 | 0.35 |
|  | Humidity + PC1 | 4 | -59.37 | 127.09 | 1.3 | 0.18 |
|  | PC1 + Maximum temperature | 4 | -59.42 | 127.19 | 1.39 | 0.17 |
|  | Tree species richness + PC1 | 4 | -59.49 | 127.32 | 1.52 | 0.16 |
|  | PC1 + PC3 | 4 | -59.67 | 127.69 | 1.9 | 0.13 |
| *Avahi occidentalis* | Tree species richness | 3 | -55.29 | 116.79 | 0.0 | 0.29 |
|  | Tree species richness + PC1 | 4 | -54.68 | 117.71 | 0.92 | 0.18 |
|  | Tree species richness + PC3 | 4 | -54.69 | 117.72 | 0.93 | 0.18 |
|  | Tree species richness + PC3 + PC1 | 5 | -54.01 | 118.54 | 1.75 | 0.12 |
|  | Tree species richness + PC2 | 4 | -55.12 | 118.59 | 1.8 | 0.12 |
|  | Humidity + Tree species richness | 4 | -55.15 | 118.64 | 1.85 | 0.11 |
| *Cheirogaleus medius* | Tree species richness | 3 | -37.2 | 80.72 | 0.0 | 0.15 |
|  | Humidity + Tree species richness | 4 | -36.16 | 80.84 | 0.13 | 0.14 |
|  | Tree species richness + PC2 | 4 | -36.38 | 81.28 | 0.56 | 0.11 |
|  | Humidity + Tree species richness + PC2 | 5 | -35.36 | 81.52 | 0.8 | 0.1 |
|  | PC2 + PC3 | 4 | -36.65 | 81.83 | 1.11 | 0.08 |
|  | PC2 | 3 | -37.79 | 81.88 | 1.17 | 0.08 |
|  | Tree species richness + PC3 | 4 | -36.69 | 81.9 | 1.19 | 0.08 |
|  | Tree species richness + PC2 + PC3 | 5 | -35.58 | 81.95 | 1.24 | 0.08 |
|  | Tree species richness + Maximum temperature | 4 | -36.85 | 82.23 | 1.51 | 0.07 |
|  | Humidity + Tree species richness + Maximum temperature | 5 | -35.82 | 82.44 | 1.73 | 0.06 |
|  | Humidity + Tree species richness + PC3 | 5 | -35.9 | 82.61 | 1.89 | 0.06 |
| *Microcebus ravelobensis* | Humidity + PC3 | 4 | -66.39 | 141.13 | 0.0 | 0.26 |
|  | Humidity + PC1 + PC3 | 5 | -65.68 | 141.88 | 0.74 | 0.18 |
|  | Humidity + PC2 + PC3 | 5 | -65.69 | 141.91 | 0.78 | 0.17 |
|  | Humidity + Tree species richness + PC3 | 5 | -65.86 | 142.25 | 1.12 | 0.15 |
|  | Humidity + PC1 + PC2 + PC3 | 6 | -64.86 | 142.45 | 1.32 | 0.13 |
|  | Humidity + PC3 + Maximum temperature | 5 | -66.08 | 142.68 | 1.55 | 0.12 |
| *Microcebus murinus* | Null | 2 | -81.84 | 167.78 | 0.0 | 0.25 |
|  | Tree species richness | 3 | -81.17 | 168.55 | 0.78 | 0.17 |
|  | Humidity | 3 | -81.5 | 169.2 | 1.42 | 0.12 |
|  | Maximum temperature | 3 | -81.5 | 169.2 | 1.43 | 0.12 |
|  | PC1 | 3 | -81.52 | 169.25 | 1.47 | 0.12 |
|  | PC2 | 3 | -81.68 | 169.56 | 1.78 | 0.1 |
|  | Humidity + Maximum temperature | 4 | -80.64 | 169.63 | 1.85 | 0.1 |
| Lemur species richness | Tree species richness + PC1 | 5 | -177.13 | 364.79 | 0.0 | 0.21 |
|  | Tree species richness | 4 | -178.27 | 364.89 | 0.1 | 0.2 |
|  | Humidity + Tree species richness | 5 | -177.69 | 365.9 | 1.11 | 0.12 |
|  | Humidity + Tree species richness + PC1 | 6 | -176.76 | 366.26 | 1.47 | 0.1 |
|  | Tree species richness + PC1 + PC2 | 6 | -176.82 | 366.38 | 1.59 | 0.1 |
|  | Tree species richness + PC1 + PC3 | 6 | -176.85 | 366.43 | 1.64 | 0.09 |
|  | Tree species richness + PC3 | 5 | -178.04 | 366.61 | 1.82 | 0.09 |
|  | Tree species richness + PC1 + Maximum temperature | 6 | -176.98 | 366.7 | 1.91 | 0.08 |

**TABLE S3** Model-averaged parameter estimates of predictors for the presence of *Eulemur fulvus*. Estimates represent the effect size of each predictor on the response variable. SE (Standard Error) indicates the precision of the estimate. The 95% Confidence Interval (95% CI) shows the range of values within which the true effect is likely to fall. Predictors are considered influential if the confidence interval does not include zero (respective results in bold). Relative Importance (RI) represents the sum of Akaike weights across all relevant models (from Table S2) in which the predictor appeared. Variables with an RI ≥ 0.8 are considered strongly supported (Burnham et al., 2011; Symonds & Moussalli, 2011).

| Species | Variables | Estimate | SE | 95% CI | Relative Importance (RI) |
| --- | --- | --- | --- | --- | --- |
| *Eulemur fulvus* | **Forest humidity** | **0.0754** | **0.0371** | **[0.0020, 0.1489]** | **1.0000** |
|  | **Tree species richness** | **0.1656** | **0.0834** | **[0.0005, 0.3308]** | **0.8419** |
|  | PC1 | 0.3269 | 0.2240 | [-0.1165, 0.7703] | 0.3934 |
|  | PC2 | 0.1060 | 0.2395 | [-0.3683, 0.5803] | 0.1198 |
|  | PC3 | 0.2446 | 0.2665 | [-0.2832, 0.7724] | 0.1645 |

**TABLE S4** Model-averaged parameter estimates of predictors for the presence of the medium-sized *Lepilemur edwardsi* and *Avahi occidentalis,* respectively. Estimates represent the effect size of each predictor on the response variable. SE (Standard Error) indicates the precision of the estimate. The 95% Confidence Interval (95% CI) shows the range of values within which the true effect is likely to fall. Predictors are considered influential if the confidence interval does not include zero (respective results in bold). Relative Importance (RI) represents the sum of Akaike weights across all relevant models (from Table S2) in which the predictor appeared. Variables with an RI ≥ 0.8 are considered strongly supported (Burnham et al., 2011; Symonds & Moussalli, 2011).

| Species | Variables | Estimate | SE | 95% CI | Relative Importance (RI) |
| --- | --- | --- | --- | --- | --- |
| *Lepilemur edwardsi* | Forest humidity | -0.0216 | 0.0233 | [-0.0677, 0.0245] | 0.1820 |
|  | Tree species richness | 0.0450 | 0.0573 | [-0.0686, 0.1585] | 0.1623 |
|  | Maximum Temperature | 0.0773 | 0.0837 | [-0.0884, 0.2430] | 0.1732 |
|  | **PC1** | **0.4378** | **0.1761** | **[0.0892, 0.7864]** | **1.0000** |
|  | PC3 | 0.1037 | 0.2094 | [-0.3111, 0.5185] | 0.1348 |
| *Avahi occidentalis* | Forest humidity | -0.0114 | 0.0211 | [-0.0532, 0.0304] | 0.1139 |
|  | **Tree species richness** | **0.1469** | **0.0611** | **[0.0261, 0.2676]** | **1.0000** |
|  | PC1 | -0.2261 | 0.2078 | [-0.6376, 0.1854] | 0.3012 |
|  | PC2 | -0.1072 | 0.1925 | [-0.4885, 0.2740] | 0.1168 |
|  | PC3 | -0.2506 | 0.2302 | [-0.7064, 0.2053] | 0.3007 |

**TABLE S5** Model-averaged parameter estimates of predictors for the presence of the small-sized *Cheirogaleus medius, Microcebus ravelobensis* and *M. murinus,* respectively. Estimates represent the effect size of each predictor on the response variable. SE (Standard Error) indicates the precision of the estimate. The 95% Confidence Interval (95% CI) shows the range of values within which the true effect is likely to fall. Predictors are considered influential if the confidence interval does not include zero (respective results in bold). Relative Importance (RI) represents the sum of Akaike weights across all relevant models (from Table S3) in which the predictor appeared. Variables with an RI ≥ 0.8 are considered strongly supported (Burnham et al., 2011; Symonds & Moussalli, 2011).

| Species | Variables | Estimate | SE | 95% CI | Relative Importance (RI) |
| --- | --- | --- | --- | --- | --- |
| *Cheirogaleus medius* | Forest humidity | 0.0376 | 0.0277 | [-0.0176, 0.0929] | 0.3522 |
|  | **Tree species richness** | **0.1452** | **0.0716** | **[0.0028, 0.2875]** | **0.8352** |
|  | Maximum temperature | 0.1135 | 0.1365 | [-0.1583, 0.3853] | 0.1298 |
|  | PC2 | -0.4016 | 0.2625 | [-0.9233, 0.1200] | 0.4508 |
|  | PC3 | -0.3487 | 0.3196 | [-0.9845, 0.2871] | 0.299 |
| *Microcebus ravelobensis* | **Forest humidity** | **0.0408** | **0.0195** | **[0.0021, 0.0795]** | **1.0000** |
|  | Tree species richness | 0.0435 | 0.0424 | [-0.0405, 0.1275] | 0.146 |
|  | Maximum temperature | -0.0516 | 0.0652 | [-0.1808, 0.0776] | 0.1175 |
|  | PC1 | 0.1696 | 0.1382 | [-0.1041, 0.4433] | 0.3080 |
|  | PC2 | -0.1894 | 0.1568 | [-0.4999, 0.1210] | 0.3051 |
|  | **PC3** | **-0.5714** | **0.1997** | **[-0.9668, -0.1760]** | **1.0000** |
| *Microcebus murinus* | Forest humidity | -0.0209 | 0.021 | [-0.0626, 0.0207] | 0.2249 |
|  | Tree species richness | 0.048 | 0.0416 | [-0.0345, 0.1304] | 0.1719 |
|  | Maximum temperature | -0.0875 | 0.0666 | [-0.2193, 0.0444] | 0.2246 |
|  | PC1 | 0.116 | 0.1454 | [-0.1719, 0.4040] | 0.1215 |
|  | PC2 | -0.0811 | 0.1429 | [-0.3642, 0.2020] | 0.104 |

**TABLE S6** Model-averaged parameter estimates of predictors for lemur species richness. Estimates represent the effect size of each predictor on the response variable. SE (Standard Error) indicates the precision of the estimate. The 95% Confidence Interval (95% CI) shows the range of values within which the true effect is likely to fall. Predictors are considered influential if the confidence interval does not include zero (respective results in bold). Relative Importance (RI) represents the sum of Akaike weights across all relevant models (from Table S3) in which the predictor appeared. Variables with an RI ≥ 0.8 are considered strongly supported (Burnham et al., 2011; Symonds & Moussalli, 2011).

| Response | Predictor | Estimate | SE | 95% CI | Relative Importance (RI) |
| --- | --- | --- | --- | --- | --- |
| *Lemur species richness* | Forest humidity | 0.0048 | 0.0049 | [-0.0049, 0.0144] | 0.2249 |
|  | **Tree species richness** | **0.0433** | **0.0140** | **[0.0157, 0.0709]** | **1.0000** |
|  | Maximum temperature | 0.0092 | 0.0165 | [-0.0235, 0.0418] | 0.0823 |
|  | PC1 | 0.0646 | 0.0422 | [-0.0189, 0.1482] | 0.5883 |
|  | PC2 | -0.0339 | 0.0427 | [-0.1184, 0.0506] | 0.0965 |
|  | PC3 | -0.0337 | 0.0472 | [-0.1272, 0.0597] | 0.1798 |

**TABLE S7** Effects of fire history and severity on forest microclimate, tree species richness, and vegetation structure (PCs 1-3). Two models (A, B) were fitted for each response variable based either on the entire dataset (model A) or only on the burnt partitions of the study sites (model B). The table reports estimates, standard errors (SE), 95% confidence intervals (CI) and the p-values with level of significance p < 5 for each predictor in all models.

| **Response Variable** | **Model** | **Predictor** | **Estimate** | **SE** | **95% CI** | **p-value** |
| --- | --- | --- | --- | --- | --- | --- |
| Forest humidity | A | Intercept | 4.36 | 0.044 | [ 4.28, 4.45] | < 0.0001*** |
|  |  | Burn status (Unburnt) | -0.00132 | 0.0242 | [-0.0487, 0.0460] | 0.984 |
|  |  | Number of fires | -0.000841 | 0.0151 | [-0.0304, 0.0287] | 0.964 |
|  | B | Intercept | 76.5 | 4.07 | [68.9, 84.4] | < 0.0001*** |
|  |  | Time since last fire | 0.170 | 0.100 | [-0.0261, 0.367] | 0.089 |
|  |  | Maximum fire severity | -2.22 | 13.6 | [-28.9, 24.4] | 0.871 |
| Tree species richness | A | Intercept | 13.7 | 1.15 | [11.4, 15.9] | < 0.0001*** |
|  |  | **Burn status (Unburnt)** | **5.05** | **1.16** | **[2.77, 7.32]** | < **0.0001***** |
|  |  | Number of fires | 0.032 | 0.5 | [-947, 1.01] | 0.948 |
|  | B | Intercept | 16.2 | 2.02 | [12.3, 20.2] | < 0.0001*** |
|  |  | Time since last fire | 0.054 | 0.068 | [-0.079, 0.188] | 0.428 |
|  |  | **Maximum fire severity** | **-33.1** | **8.25** | **[-49.2, -16.9]** | < **0.0001***** |
| PC1 | A | Intercept | 0.162 | 0.384 | [-0.591, 0.916] | 0.670 |
|  |  | **Burn status (Unburnt)** | **1.11** | **0.328** | **[0.464, 1.75]** | < **0.001**** |
|  |  | Number of fires | -0.254 | 0.135 | [-0.520, 0.0105] | 0.059 |
|  | B | Intercept | -0.343 | 0.387 | [-1.10, 0.415] | 0.376 |
|  |  | **Time since last fire** | **0.0510** | **0.0135** | **[0.0246, 0.0775]** | **0.0001***** |
|  |  | **Maximum fire severity** | **-10.5** | **2.62** | **[-15.6, -5.39]** | **< 0.0001***** |
| PC3 | A | Intercept | -0.0782 | 0.2610 | [-0.590, 0.433] | 0.765 |
|  |  | Burn status (Unburnt) | 0.0802 | 0.295 | [-0.498, 0.659] | 0.786 |
|  |  | Number of fires | 0.0379 | 0.113 | [-0.184,0.260] | 0.738 |
|  | B | Intercept | -0.665 | 0.394 | [-1.44, 0.107] | 0.091 |
|  |  | **Time since last fire** | **0.0244** | **0.0113** | **[0.00230, 0.0465]** | **0.030*** |
|  |  | Maximum fire severity | 3.02 | 2.13 | [-1.16, 7.20] | 0.157 |

**Table S8** Result of the Post-hoc Tukey test (R package emmeans, Lenth, 2023) used to compare the three forest zones (Burnt/Transition/Unburnt) regarding their forest humidity, tree species richness, PC1 and PC3*.* In bold: significant results.

|  | Models | Pairwise comparisons | Estimate | Standard error | Z-ratio | P-value |
| --- | --- | --- | --- | --- | --- | --- |
|  | Forest humidity | Burnt-Transition | 1.688 | 1.81 | 0.932 | 0.6211 |
|  |  | Burnt-Unburnt | 0.506 | 1.58 | 0.321 | 0.9447 |
|  |  | Transition-Unburnt | -1.181 | 2.32 | -0.510 | 0.8669 |
|  | Tree species richness | Burnt-Transition | 0.077 | 0.084 | 0.915 | 0.6305 |
|  |  | Burnt-Unburnt | -0.330 | 0.083 | -3.966 | **< 0.001**** |
|  |  | Transition-Unburnt | -0.408 | 0.110 | -3.705 | **< 0.001**** |
| PC1 | | Burnt-Transition | -0.212 | 0.327 | -0.649 | 0.7933 |
|  |  | Burnt-Unburnt | 1.134 | 0.342 | 3.311 | **< 0.01*** |
|  |  | Transition-Unburnt | 1.346 | 0.424 | 3.178 | **< 0.01*** |
| PC3 | | Burnt-Transition | -0.194 | 0.374 | -0.694 | 0.7676 |
|  |  | Burnt-Unburnt | 0.05 | 0.298 | 0.182 | 0.981 |
|  |  | Transition-Unburnt | 0.249 | 0.374 | 0.665 | 0.7840 |

**References**

Burnham, K. P., Anderson, D. R., & Huyvaert, K. P. (2011). AIC model selection and multimodel inference in behavioral ecology: some background, observations, and comparisons. *Behavioral Ecology and Sociobiology*, *65*(1), 23–35. <https://doi.org/10.1007/s00265-010-1029-6>.

Lenth, R. (2023). emmeans: Estimated Marginal Means, aka Least-Squares Means_. R package version 1.8. 5.

Symonds, M. R. E., & Moussalli, A. (2011). A brief guide to model selection, multimodel inference and model averaging in behavioural ecology using Akaike’s information criterion. *Behavioral Ecology and Sociobiology*, 65(1), 13–21. https://doi.org/10.1007/s00265-010-1037-6.
